# Spatial self-organization of cancer stem cell niches revealed by live single-cell imaging

**DOI:** 10.1101/2024.06.07.597918

**Authors:** Mathilde Brulé, Anaïs Horochowska, Emeline Fontaine, Raoul Torero-Ibad, Flavie Woesteland, Marie Denoulte, Jean Pesez, Eric Adriaenssens, Robert-Alain Toillon, Xuefen Le Bourhis, Benjamin Pfeuty, Chann Lagadec, François Anquez

**Author notes:** These authors contributed equally to this work (Mathilde Brulé, Anaïs Horochowska). These authors also contributed equally to this work (Chann Lagadec, François Anquez). Nantes Université, INSERM, CNRS, Université d’Angers, CRCI2NA, Nantes, France.

## Abstract

Phenotypic plasticity is a major factor of tumor heterogeneity and treatment resistance. In particular, cancer stem cells (CSCs) represent a small subpopulation within tumors with self-renewal and tumor-forming capabilities. Understanding reprogramming, maintenance, and lineage properties of CSCs requires dedicated tools to disentangle the respective influences of phenotypic inheritance and cell-cell interactions. Here we set up ultra-wide field microscopy of breast cancer cell lines expressing a stemness fluorescent reporter for several days. The fluorescent reporter distinguishes three phenotypes: cancer stem cells (CSCs), cancer differentiated cells (CDCs) and intermediate/transiting cancer cells (iCCs). Spatial statistics indicate significant zonation, aka phenotypic niches, with CSC clustering near each other but away from CDCs. Surprisingly, single cell time series reveal spontaneous reprogramming events from CDC to CSC even in unperturbed populations. We identify that such transitions are prone to arise during the cell cycle. Moreover, lineage analysis shows that the phenotype is partially inherited from ancestor cells. However, such heredity is not sufficient to explain the spatial properties of the cell population, which also depend on cell-cell interactions. Indeed, we identified that phenotypic transitions of cancer cells are influenced by the phenotypic state of neighboring cells. Reprogramming into CSCs is respectively promoted and inhibited by the presence of CSCs and CDCs in the neighborhood. Altogether, our results disentangle how phenotypic inheritance and intercellular interactions orchestrate the spatio-temporal self-organization of cancer cell heterogeneity, maintaining a subpopulation of CSCs within niches.

## Introduction

Tumors constitute a diverse array of cells, encompassing transformed cancer cells, supportive cells, and cells infiltrating the tumor. Tumor heterogeneity extends beyond malignant cells alone, as tumors constitute intricate ecosystems housing various cell types such as endothelial cells, macrophage and lymphocyte cells, cancer-associated fibroblasts, and an intricate extracellular matrix network, contributing to spatial and temporal disparities in the tumor microenvironment (Junttila and De Sauvage, 2013; Lu et al., 2012). Moreover, these diversities result in variations among patients, but also within individual tumors, influencing the spatial and temporal characteristics of tumors. This variability significantly impacts how tumors respond to drugs and ultimately affects the outcome of the disease (Kashyap et al., 2022).

Throughout the cancer cell population, heterogeneity arises from mutation and epigenetic origins, but also from the regulation of different phenotypes within the cancer cell population. The clonal evolution model postulates that stochastic mutations in individual tumor cells provide a basis for adaptation and selection of the fittest tumor clones, driving intra-tumor heterogeneity through selection. Clones with advantageous traits proliferate, while those less fit are out-competed and may eventually disappear. Importantly, these advantages may vary over time and with surrounded environment, different areas of the tumor favoring distinct clone types based on environmental conditions (Anderson et al., 2006; Sottoriva et al., 2010; Waclaw et al., 2015). While initial observations indicated the presence of sub-clones within tumors exhibiting differences in genetic makeup and response to chemotherapy (Shapiro et al., 1981; Yung et al., 1982), recent profiling efforts utilizing comprehensive sequencing and methylation analysis across various tumor regions have unveiled multiple distinct clones harboring unique genetic mutations and epigenetic patterns within a single tumor (Anderson et al., 2011; Gerlinger et al., 2012). Meanwhile, the cancer stem cell (CSC) model suggests that only a subset of cancer cells possess the capacity for indefinite self-renewal, giving rise to progenitors and differentiated cells which sustain tumor growth in a hierarchical manner akin to normal tissue hierarchy maintained by healthy normal stem cells. Consequently, CSCs generate cellular diversity by establishing a differentiation hierarchy within the tumor (Colacino et al., 2018). However, this hierarchy is not strictly unidirectional, as terminally differentiated cells can revert to a stem cell-like state under specific conditions, a phenomenon known as cell plasticity (Friedmann-Morvinski and Verma, 2014; Wahl and Spike, 2017; Brown et al., 2022). The concept of cell plasticity reconciles elements of both stochastic and CSC models, whereby mutations in differentiated cells can confer self-renewal ability, establishing new hierarchical CSC clones and increasing functional diversity within the tumor (Wahl and Spike, 2017). The capacity of cells to transition between states through various programs, such as Epithelial-Mesenchymal Transition (EMT), suggests that CSCs may not always be predetermined; instead, stemness can be considered as a cellular state that can be gained or lost reversibly. In essence, cellular plasticity enables dynamic transitions between CSCs and non-CSCs (Chaffer et al., 2011; Gupta et al., 2019). Furthermore, distinct subsets of CSCs may occupy different positions along the epithelial-mesenchymal axis and have the potential to undergo inter-conversion (Liu et al., 2014; Bocci et al., 2018, 2019a).

Upon homeostatic conditions, cancer cells exhibit stable populations of both stem-like and differentiated cells. The activity of CSCs is orchestrated by a multitude of pluripotent transcription factors including OCT4, SOX2, NANOG, KLF4, and MYC (Zhang and Wang, 2008). Furthermore, numerous intracellular signaling pathways play pivotal roles in regulating CSC behavior. These pathways include Wnt (Kanwar et al., 2010), Notch (Wang et al., 2010), Sonic Hedgehog (Li et al., 2007), NF-κB, JAK-STAT, PI3K/AKT/mTOR, TGF/SMAD, and PPAR (Yang et al., 2020). In addition to these intracellular pathways, various extracellular influences are also crucial. These influences include vascular niches, hypoxia, tumor-associated macrophages, cancer-associated fibroblasts, cancer-associated mesenchymal stem cells, extracellular matrix, and exosomes (Plaks et al., 2015; Yoshida and Saya, 2016; Oshimori et al., 2021). Both paracrine and juxtacrine mediated signalling spatially shape the tumor leading to the formation of CSC niches. Interestingly a complex equilibrium maintains a balance between CSCs and differentiated cancer cells. As an example, differentiated cancer cells secrete factors like Interleukin-6 (IL6) (Liu et al., 2015) and BDNF-NTRK2-VGF (Wang et al., 2018), NGF/p75NTR axis (Tomellini et al., 2015), fostering the survival and self-renewal of CSCs in breast cancer and glioblastoma. While CSCs secrete factors like DKK1 pushing forward differentiation (Wu et al., 2022). Perturbing this equilibrium results in unbalancing the system and induces phenotypic switch. Indeed, phenotypic plasticity of CSC, also called reprogramming from non-CSC into CSC, was first observed upon perturbations such as anticancer treatments ((Lagadec et al., 2012)) or specific environmental condition as hypoxia (Li et al., 2009). After treatment, enriched Breast Cancer Stem Cells (BCSCs) swiftly restore the parental cell composition, indicating the inclination of BCSCs to differentiate under such circumstances (Bidan et al., 2019; Gupta et al., 2011; Iliopoulos et al., 2011).

In view of these numerous pathways involved in phenotypic balance, in order to propose efficient therapeutic strategies, there is a need for *ex vivo* assays emulating the spatial arrangement of cell types in tumors, as can be done to recapitulate essential steps of development such as germ layer and axial patterning (Warmflash et al., 2014). However, the diversity of phenotypes and interaction motifs and signalling pathways involved make it difficult to understand how CSCs niche auto-organizes in space and time. Some general insights about phenotypic heterogeneity and spatial organization of stem cell niche could be gained through system-level approaches. Indeed, regardless the signaling pathways at play, the main sources of spatial self-organization of cell phenotypes are fundamentally influenced by a few cell-fate processes, namely differentiation, division, motility and death (Kicheva et al., 2012; Grace and Hütt, 2015; Landge et al., 2020). While the complexity of tumor heterogeneity arises from the diversity of intracellular and extracellular signaling mechanisms that couple all these processes, some key determinants can be unveiled by focusing on the initial and spontaneous trends to self-organize in unperturbed environments. We therefore hypothesize that cultured cancer cell lines show some rudimentary forms of self-organization featured with a spatial segregation of cancer stem cell niche.

Live cell imaging tools provide most spatio-temporal information sufficient to track the main processes involved in spatial self-organization. This wealth of data facilitates a system-level approach to investigate the intricate interplay between the niche effect and the homeostatic capacity of tumor models. It also aids in identifying key factors that could disrupt the cancer stem cell niche and cellular plasticity. In this work, we investigate the interplay between the temporal transitions and the spatial distribution of stemness phenotypes in population of breast cancer cell upon unperturbed and steady culture conditions. To this aim, we developed an ultra-wide field imaging system capable of tracking thousands of single-cells in space and time for days. Using mNeptune fluorescent protein expression under the control of the promoter of ALDH1A1 (pALDH1A1:mNeptune) as a stemness reporter for breast cancer cells (Bidan et al., 2019), we conducted a series of analyses to characterize the spatial and temporal features of phenotypic heterogeneity. From the broad distribution of fluorescent marker over the population, statistical analysis identifies two main phenotypes associated to stemness and differentiation traits, as well as an intermediate state. Point pattern analysis uncovered spatial segregation between stem-like and differentiated phenotypes. Notably, we observed phenotypic transitions, including reprogramming events from CDCs to CSCs phenotypes, even within unperturbed cancer cell populations. Analysis of cell fate further underscored the significant role of intercellular signaling in phenotypic transitions and CSC reprogramming. Altogether, our findings demonstrate that cancer stem cell niche spontaneously emerge in unconstrained population of cancer cells through the interplay of phenotypic inheritance across generations and intercellular communications that stabilize the stemness phenotype.

## Results

### Ultra-wide field imaging characterizes cancer cell phenotypic diversity and its spatial distribution

To distinguish CSCs from non-CSCs and to track the spatio-temporal dynamics of cells, we previously developed a reporter based on mNeptune fluorescent protein expression under the control of the promoter of ALDH1A1 (pALDH1A1:mNeptune) (Bidan et al., 2019). In 2D cell culture conditions, CSC frequency is rather low (from <1% up to 5%) (Fillmore and Kuperwasser, 2008). Thus, we set up ultra-wide field microscopy to investigate the phenotypic diversity of breast cancer cells with sufficient statistics.

SUM159-PT and MDA-MB-231 cells were imaged with a high Numerical Aperture objective to allow detection of the faint pALDH1a1:mNeptune signal. A 6.8mm × 6.8 mm wide field of view is reconstructed by assembling 60X high resolution images (***Figure 1***). Images reveal a large diversity in single cell mNeptune fluorescence intensity, with majority of cells exhibiting no or low fluorescence and a small fraction of cells expressing high fluorescence (Figure 1C). This observation qualitatively aligns with a low CSC fraction reported in breast cancer cells (Fillmore and Kuperwasser, 2008). At larger scale (Figure 1A), reconstructed images indicate high spatial heterogeneity with small regions populated by cells exhibiting high fluorescence.

**Figure 1.**
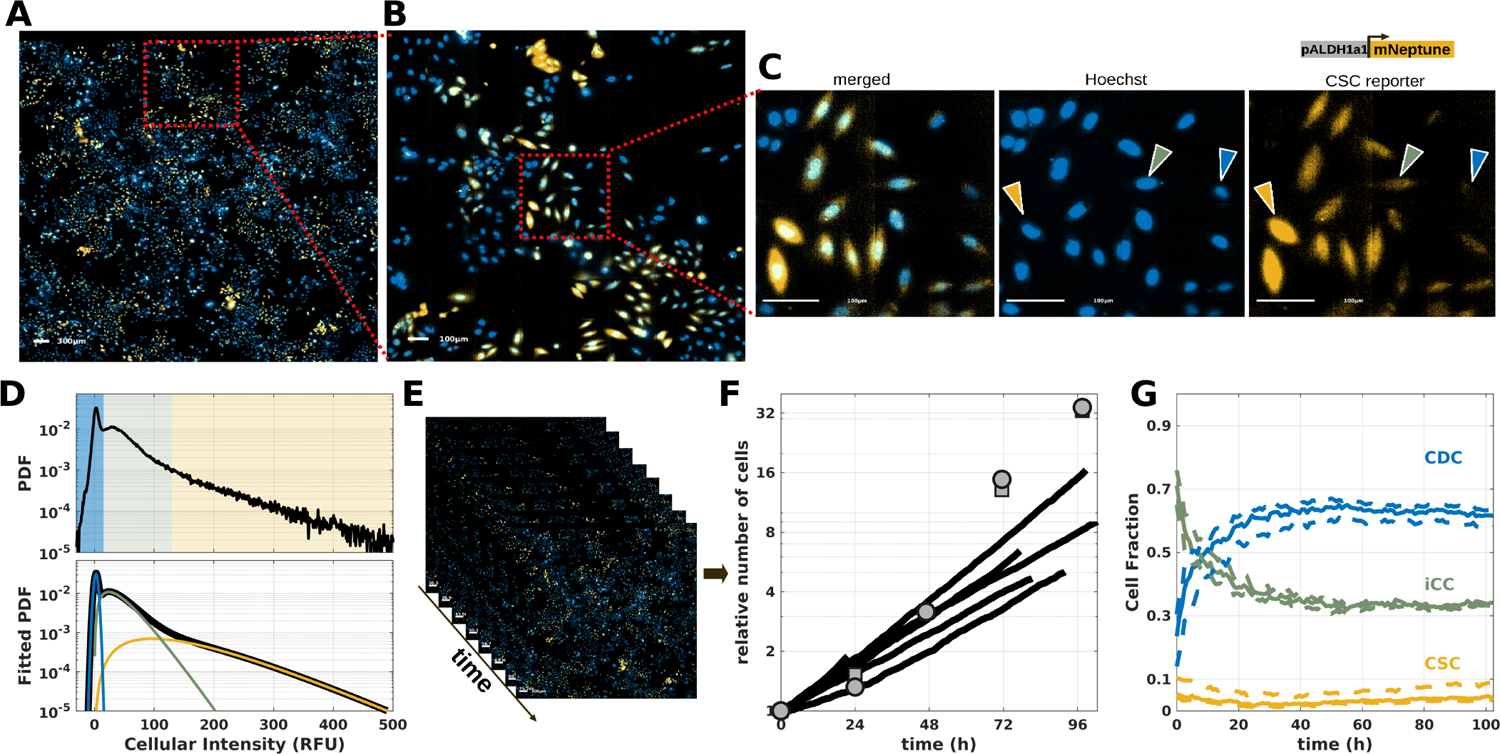
Ultra-wide field time-lapse imaging of CSC fluorescent reporter with single cell resolution. (**A**-**C**) Representative epi-fluorescence images of SUM159-PT breast cancer cells stably transfected with the CSC reporter, pALDH1a1:mNeptune (Bidan et al., 2019). Hoechst (blue). CSC reporter (yellow). (**A**) Reconstructed ∼6.8mm × ∼6.8 mm field of view 72h after the beginning for a representative experiment. The full field of view is reconstructed by assembling 32 × 32 higher resolution images. Scale bar 300*μm*. (**B**) A 4X zoom on a subset of the same image. Scale bar 100*μm*. (**C**) A 16X zoom on a subset of the full width image. CSC reporter signal only (right), Hoechst signal only (middle) and merged Hoechst and CSC reporter (left). Scale bar 100*μm*. The yellow, green and blue arrows respectively indicate high, medium and low fluorescent cells. (**D**) Upper panel: probability density function of single cell CSC reporter signal (extracted by means of image processing, see material and methods) for one representative experiment. Lower panel: fit of the PDF with a the sum of three probability distributions (black line, see material and methods). A normal distribution for cells with low intensities (blue, mean *μ* ≈ 2.6 and standard deviation *σ* ≈ 3.3) and a gamma distribution for cells with intermediate intensities (green, shape parameter *k* ≈ 2.3 and scale parameter *k* ≈ 18.2) and another gamma distribution for cells with higher fluorescence level (yellow, shape parameter *k*_+_ ≈ 2.7 and scale parameter *k*_+_ ≈ 55.7). For all experiments, we define two fluorescence thresholds (*θ*_−_ = 15.5 and *θ*_+_ = 129.5) to separate the three cell populations (see material and methods): cells with signal lower than *θ*_−_ are considered as Differentiated Cancer Cells (CDCs); cells with signal higher than *θ*_+_ are considered as Cancer Stem Cells (CSCs); and cells with signal between than *θ*_−_ and *θ*_+_ are labelled as intermediate fluorescence Cancer Cells (iCCs). (**E**) Time-resolved analysis *via* time-lapse imaging where a full field of view (∼6.8mm × ∼6.8 mm) is acquired every ∼ 45 min. (**F**) Time evolution of the relative cell number for 5 different experiments under the microscope (black lines) or grown in standard conditions with (gray squares) or without (gray circles) Hoechst staining. The growth curve is fitted (not shown) with an exponential function and the mean doubling time is between 27 and 36h. (**G**) Time evolution of the cell fraction of the three phenotypes (CDC blue line, iCC green line and CSC yellow line) defined by the intensity thresholds *θ*_−_ and *θ*_+_. Dashed lines shows a sensitivity to thresholds (*θ*_−_ and *θ*_+_) analysis. Time evolution of proportions of each phenotype is shown in dashed lines for 12.5≤ *θ*_−_ ≤17.5 and 84.5≤ *θ*_+_ ≤148.5. Figure 1**—video 1.** Representative time-lapse of SUM159-PT breast cancer cells stably transfected with the CSC reporter Figure 1—figure supplement 1. Repeatability of fluorescence distribution and time evolution in SUM159-PT cells Figure 1—figure supplement 1. Repeatability of fluorescence distribution and time-evolution in MDA-MB-231 cells

To further quantify distribution of SUM159-PT sub-population, in addition to the CSC reporter, we used cell segmentation and image processing (see material and methods). At cellular level, the CSC reporter signal is evenly spread over the cell volume (Figure 1C). Thus, we used average nuclear mNeptune fluorescence as a proxy for single cell phenotype. We extracted single cell fluorescence for up to 10^4^ individual cells per reconstructed field of view. We observed that single cell fluorescence show a tailed distribution (Figure 1D and ***Figure 1***—figure Supplement 1 A) with ∼ 95% of cells with fluorescence below 130 Relative Fluorescence Units (RFU). More precisely the empirical fluorescence distribution is well described by the sum of three probability density functions (lower panel of Figure 1D). A normal distribution centered at low fluorescence level (2.6 RFU) likely corresponds to cancer differentiated cells (CDCs) while, on the other side, a gamma distribution with a mean of ∼ 150 RFU corresponds to CSCs. Additionally, a third distribution, also fitted by a gamma distribution (mean ∼ 42 RFU) indicates the presence of intermediate cancer cells (iCCs) with characteristics that presumably lie between CDCs and CSCs. The cell fluorescence distribution is reproducible from one experiment to another, although weights of each sub-population may slightly vary (***Figure 1***—figure Supplement 1A). Similar results are obtained with another cell line, MDA-MB-231 (figure Supplement 2A), for which the fluorescence distribution is also captured by the sum of three distinct probability density functions centered on low, intermediate and high fluorescence intensity. The intersection between the fitted probability density functions (PDF) defines thresholds that are further used to approximate the fraction of cells in the respective phenotypic states, CDC, iCC, or CSC.

Finally, our setup allows time-lapse imaging of live cells (Figure 1E and***Figure 1***—video 1). Unperturbed SUM159-PT cells are tracked for up to 5 days at an acquisition rate of one frame every 45 min. Analyzing cell growth over time, we find that the growth rate is similar under our imaging system compared to cells grown in a standard incubator (Figure 1F). We also do not observed qualitative morphological differences between cells growing under imaging and in standard conditions. This indicates that imaging with our top stage incubator does note introduce major side effects one th cancer cell population. By means of cell segmentation and image processing, we recovered the time evolution of the three identified populations (Figure 1G). After a small drop during the first 30h, the fraction of CSCs remains constant (∼ 5%) for the entire duration of the experiment. This observation is consistent with previous reports in breast cancer cells with our CSC reporter (Bidan et al., 2019) or using other CSC markers (Lagadec et al., 2012). On the other hand, CDC and iCC sub-populations are much more dynamic during a transient phase that last approximately 24 to 36h after which their proportions stabilize to ∼ 65% for CDCs and ∼ 30% for iCCs. This behaviour is reproducible from one experiment to another (figure Supplement 1B). And we observe a similar trend with the other cell line, MDA-MB-231 (figure Supplement 2B).

Interestingly, we do not find a significant difference between the mean division rates (0.02 to 0.04ℎ^−^1) of all three populations with SUM159-PT (figure Supplement 1C) and MDA-MB-231 (figure Supplement 2C) cell lines. Specifically, the division rate of CDCs does not significantly exceed that of iCCs during the transient phase. This suggests that the progressive establishment of steady proportion of cancer cell phenotypes relies on dynamics balance between cell-fate transition events, initially dominated by transitions from intermediate to differentiated phenotypes.

### Point pattern analysis unveils spatial segregation of phenotypes with clusters of CSCs

To quantitatively characterize spatial heterogeneity, we employed functions from point pattern analysis (Cressie, 2015), a methodology recently applied to biological systems (Parra, 2021). Each cell is characterized by both its spatial coordinates (X,Y) and a specific phenotype (CDC, iCC or CSC (Figure 2A left panel). We estimated the empirical Point Correlation Function (PCF), denoted as *g*(*r*), which is a function of a distance, *r*. The value of *g*(*r*) is computed by counting events within a thin annulus of radius *r* with a thickness of 5 μm (see Material and Methods). PCF measures the increase or decrease of the likelihood of finding an event at a distance *r* compared to what would be expected under complete spatial randomness (CSR). *g*(*r*) greater than 1 indicates a more clustered pattern than CSR while a value smaller than 1 indicates a more dispersed pattern than CSR (Figure 2A, right panels).

**Figure 2.**
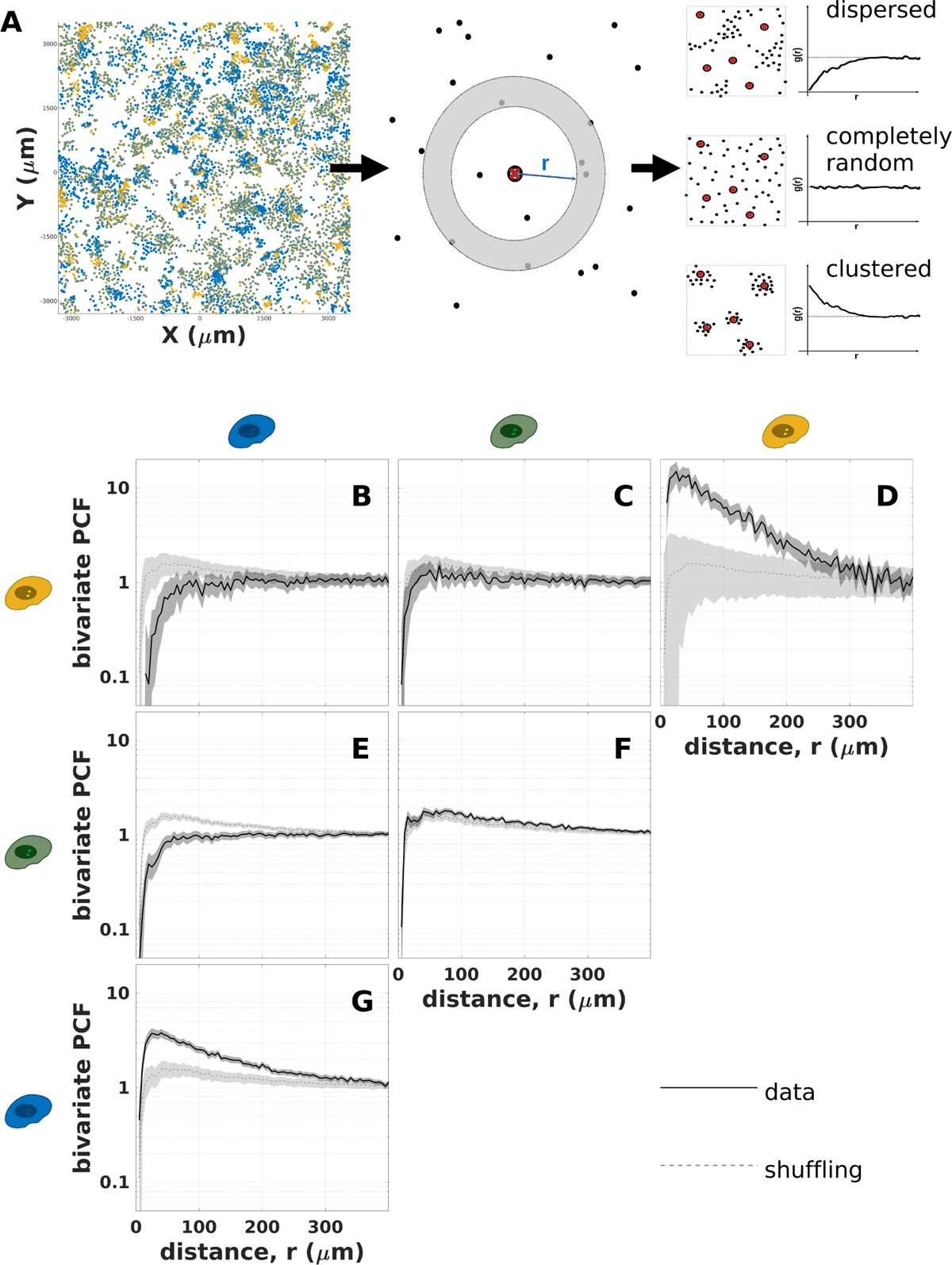
Statistical analysis of the spatial distribution of SUM159-PT phenotypes. (**A**) Workflow and interpretation of Point Pattern Analysis (see Material and Methods for details). For a given experiment and time point (here 90h), detected cells are characterized by their spatial coordinates (X,Y) and phenotypes (CDC in blue, iCC in green or CSC in yellow). The corresponding Point Correlation Function (PCF, **g**(**r**)) is computed. PCF measures the increase or the decrease of the likelihood of finding an event at a distance **r** compared to what would be expected under complete spatial randomness (CSR). **g**(**r**) greater than 1 indicates a more clustered pattern than CSR while a value smaller than 1 indicates a more dispersed pattern than CSR. (**B**-**G**) Bivariate PCF, **g*_*xy*_*(**r**) = **g*_*yx*_*(**r**), for a representative experiment 90h after beginning of imaging where **x** and **y** represent cell phenotypes (CDC, iCC or CSC). The dark lines are the measured PCFs (data). The gray shaded area is the 99% confidence interval obtained from bootstrap resampling (sensitivity analysis, see Material and Methods). The dashed gray lines are the PCF obtained from phenotype shuffling (control). The gray shaded area is the 99% confidence interval obtained from 1500 repeats of shuffling. (**B**) Bivariate PCF measuring spatial correlation between CSC and CDC. (**C**) Bivariate PCF measuring spatial correlation between CSC and iCC. (**D**) Bivariate PCF measuring spatial auto-correlation of CSC. (**E**) Bivariate PCF measuring spatial correlation between iCC and CDC. (**F**) Bivariate PCF measuring spatial auto-correlation of iCC. (**G**) Bivariate PCF measuring spatial auto-correlation of CDC. Figure 2—figure supplement 2. Univariate (regardless of phenotype) Point Correlation Function of SUM159PT cells Figure 2—figure supplement 2. Time evolution of bivariate Point Correlation Functions of SUM159-PT cells Figure 2—figure supplement 2. Repeatability of bivariate Point Correlation Function of SUM159-PT cells Figure 2—figure supplement 2. Repeatability of bivariate Point Correlation Function of MDA-MB-231 cells

We first computed the univariate PCF, for which cell phenotypes are not distinguished (figure Supplement 1). At very short distances (*r* ≤ 10 μm), the univariate PCF remains below 1, indicating that cells do not overlap in our experimental conditions. Subsequently, the function peaks at approximately 1.5 to 2 for radii of around 40 to 60 μm (depending on the experiment), reflecting the close proximity of sister cells and the formation of micro-colonies. At larger scales, the PCF gradually approaches 1, indicating a complete spatial randomness.

To investigate the spatial correlation between phenotypes, we computed the bivariate PCF. For each phenotype, we screened the presence of the same or other phenotypes within its surroundings. Because the univariate PCF has a well defined shape reflecting tissue structure, we needed to disentangle properties due to putative relationship between phenotypes from the univariate spatial organization. To achieve this, any statistical test must account for the sample size to correctly estimate the confidence interval (Cressie, 2015). We thus compared the experimental bivariate PCF with the one obtained from phenotype shuffling in which cell fluorescence are randomly permuted (see materials and methods). Overall, no significant differences between shuffing and bivariate PCF are found for iCC versus iCC or CSC (Figure 2C and F). In contrast, CSCs (respectively CDCs) are more clustered with CSCs (respectively CDCs) compared with shuffling or complete spatial randomness (CSR) (Figure 2D and G). This effect is more pronounced with CSCs for which *g*_++_(*r* ≈ 15 μm) is about 20 times greater than CSR and 10 times greater than shuffling (Figure 2D). Fitting *g*_++_(*r*) to an exponential curve indicated a characteristic length of ∼ 50 μm at the beginning of the experiment slowly increasing to ∼ 100 μm after 4 days (***Figure 2***—figure Supplement 2). Finally, at short range (*r* ∼ 15 μm) both CSC and iCC are excluded from regions with a high density of CDC (Figure 2B and E). Again, the effect is more pronounced with CSC versus CDC for which *g*_+−_(*r* ≈ 15 μm) is found to ∼ 10 times smaller than CSR and 15 times smaller than shuffling (Figure 2B). This clustered pattern is observed at all time points (figure Supplement 2) and is reproducible from one experiment to another (figure Supplement 3). We also applied point pattern analysis to data collected for the other breast cancer cell line, MDA-MB-231, and similar results are found, though with weaker correlation (figure Supplement 4).

In summary, we observe a spatial patterning of cancer cell phenotypes, where cells given phenotype (low or high fluorescence intensities) tend to cluster with cells with the same phenotype. Notably, CSCs tend to form clusters of averaged diameter of 100 μm (about 10 cell width). Intermediate phenotypes does not show significant spatial correlation with other CSCs nor iCCs, but display negative correlation with CDCs, thereby suggesting that iCCs are likely to transit between CDCs and CSCs at the interface of CSC-dense and CDC-dense areas.

### Machine learning applied to single-cell time series identifies phenotypic transitions such as CSC reprogramming events

Because proliferation rate is similar for all phenotypes, the robust establishment of a specific proportion of different phenotypes throughout the population is expected to solely rely on dynamic transitions between stem, intermediate and differentiated states. It is questionable whether these transitions mainly occur during the initial transient phase to establish a steady proportion or throughout the entire course of steady population growth. To address these issues, we used unsupervised partitioning analysis of single-cell time series (SCTS) to, without prior knowledge of time scales or specific shapes, identify distinctive temporal patterns of differentiation or reprogramming events. Automated tracking of cell trajectories, detection of division events and single-cell fluorescence extracted by means of image processing (see materials and methods) allow us to reconstruct single-cell time series (SCTS). Then, as depicted on Figure 3A, cells for which the full cell cycle could be monitored were selected for further analysis. For SUM159-PT cells, we collected 19,620 SCTS from 5 different experiments. We used cell-cycle progression (time relative to the cell cycle duration) as a metric for time evolution and each time series is re-sampled so that all SCTS share the same number of points along cell cycle progression. Euclidean distance is used as a di-similarity measurement and k-medoids (Kaufman, 1990) is used to partition SCTS and the GAP statistics (Tibshirani et al., 2001) is used to determine the relevant number of clusters. This procedure results in 16 clusters. However the last cluster (#16) contains SCTS from only 2 experiments, while all other clusters cover all experiments (figure Supplement 1A and B). This last cluster was excluded from further analysis and we finally considered 15 clusters (Figure 3) with 99.9% of the SCTS clustered. The resulting clustering did not show significant bias for any experiment (figure Supplement 1A and B).

**Figure 3.**
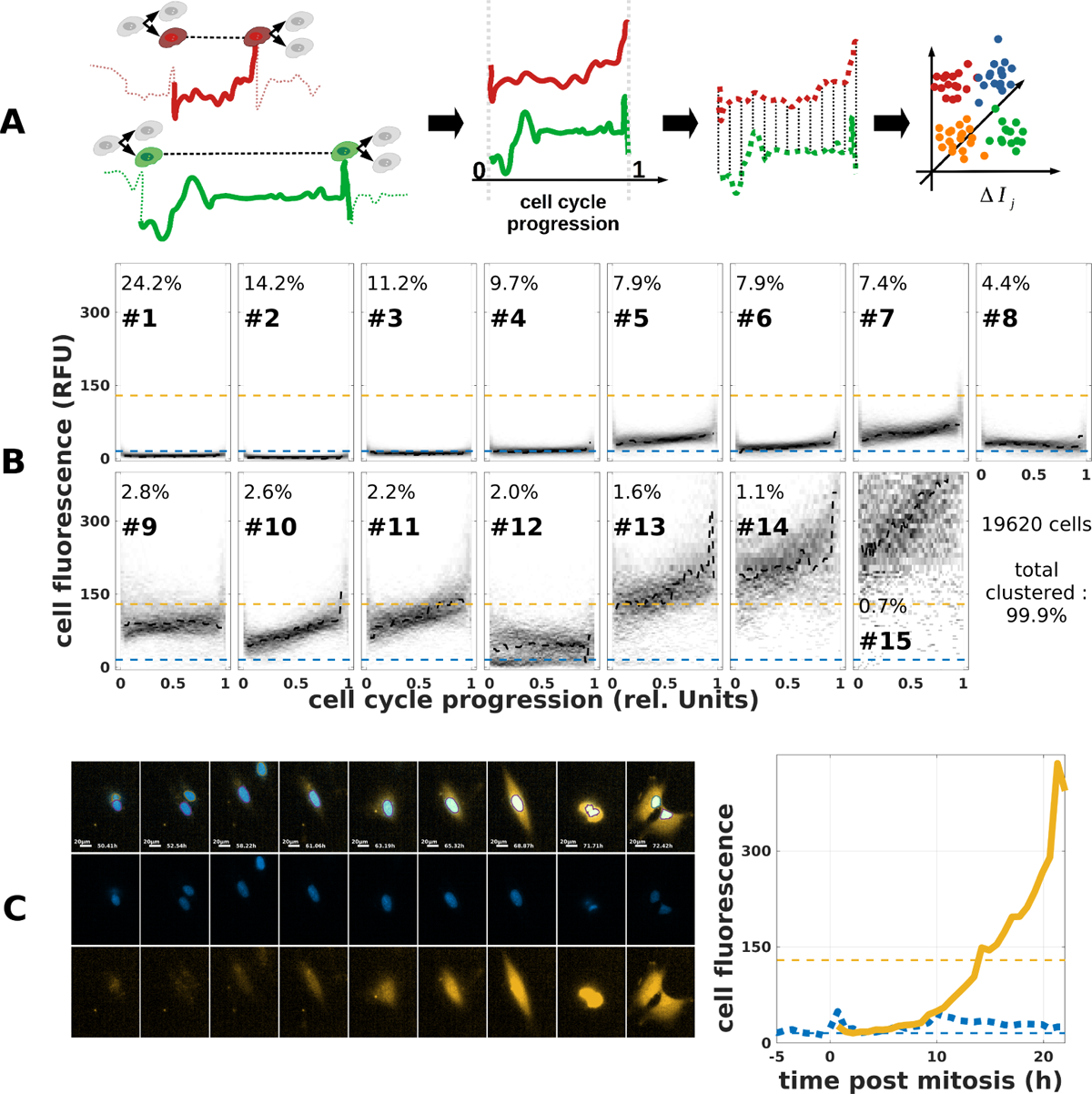
Fluorescent signal dynamics at single cell level. (**A**) **Schematic representation of single cell time-series analysis:** Selection of cells for which signals is recorded throughout a whole cell cycle. Time series are (*i*) resampled to 60 points using cell cycle progression as a time metric for time, (*ii*) compared using a Euclidean distance, and (*iii*) partitioned around 16 medoids. (**B**) **Results of single cell time-series clustering for SUM159PT cells:** Clusters are numbered by increasing size. The percentage of cells within each cluster is indicated. Are shown scatter plot of each time-series within each cluster. (**C**) Example of a cell with a significant increase of fluorescent signal associated to de-differentiation into CSC. Left panel: time-lapse images (scale bar 20*μm*). bottom: m-Neptune fluorescence; middle Hoechst and top: merged. Right panel: Time evolution of fluorescence signal of the cell (yellow trace) and the one of its mother cell (blue dotted) and sister cell (blue dashed). Figure 3—figure supplement 3. Meta-analysis of identified SCTS clusters Figure 3—figure supplement 3. Transitionning and non-transitionning SCTS clusters in SUM159PT cells Figure 3—figure supplement 3. Transitionning and non-transitionning SCTS clusters in MDA-MB-231 cells Figure 3—figure supplement 3. SCTS across generations in SUM159PT cells

Partitioning analysis effectively captures the phenotypic diversity as the procedure results in SCTS categorized based on their average fluorescence signal (Figure 3B). More generally, clustered SCTS could be classified into two main categories: approximately 70% of cells exhibiting a stable signal (figure Supplement 2B), while approximately 30% of the cells within a transiting state undergoing an increase toward higher fluorescence (figure Supplement 2A). It is important to note that these transitions occur gradually throughout the cell cycle, without any abrupt changes. We obtained similar results for the other cell line MDA-MB-231 (figure Supplement 3) for which we found approximately 50% transitioning SCTS. However we note that transitioning SCTS clusters of MDA-MB-231 also include SCTS with decreasing fluorescence ((figure Supplement 3 clusters #14, 7 and 8).

Notably, the partitioning analysis unveils clusters (#10 and 11) with ascending signal and for which cells transition from a fluorescence level close to the CDC threshold to a fluorescence level close or above CSC threshold (Figure 3B and figure Supplement 2A). Clusters #10 and 11 comprise 4.8% of the analyzed time series (Figure 3B) and are not limited the transient phase at the beginning of the experiment (figure Supplement 1 C). This last observation support the idea that steady proportion of phenotypes arises from dynamic conversions throughout experiment time course. More specifically, among cells within ascending clusters, we found representative instances of single cell transitioning from CDC to CSC (Figure 3C). Interestingly, progenitors from the same mother cell can have very different fates. Indeed, the sister of the reprogramming cell depicted in Figure 3C does not exhibit transition nor significant fluorescence variation (Figure 3C left panel).

To qualitatively visualize whether phenotype is transmitted to the cell progeny, we plot the clustered SCTS together with the mother SCTS and daughters SCTS when available (figure Supplement 4). While the fluorescence level of a mother seems to be preserved for its daughter, cell division nevertheless introduces some variability, which will be studied more systematically in next section. Remarkably, transiting SCTS are not limited to the transient phase (0 to 24/36h) for which we observed exchange between iCC and CDC (Figure 1G). These reprogramming SCTS are also detected during the second phase of the experiment when steady state is already reached at the population level (figure Supplement 1C). This strongly suggests that a steady proportion of CSC fraction is maintained through a dynamic equilibrium between reprogramming and differentiating trends.

### Lineage analysis reveals phenotypic inheritance through symmetric division

To investigate how cell cycle influences phenotypic changes, we examined the fluorescence variations across generations. Figure 4A shows an example of lineage reconstruction with a depth of three generations. From a mother cell, we kept track of its 2 daughters which gave rise to 4 grand-daughters for which we could detect 6 out of 8 great grand-daughters.

**Figure 4.**
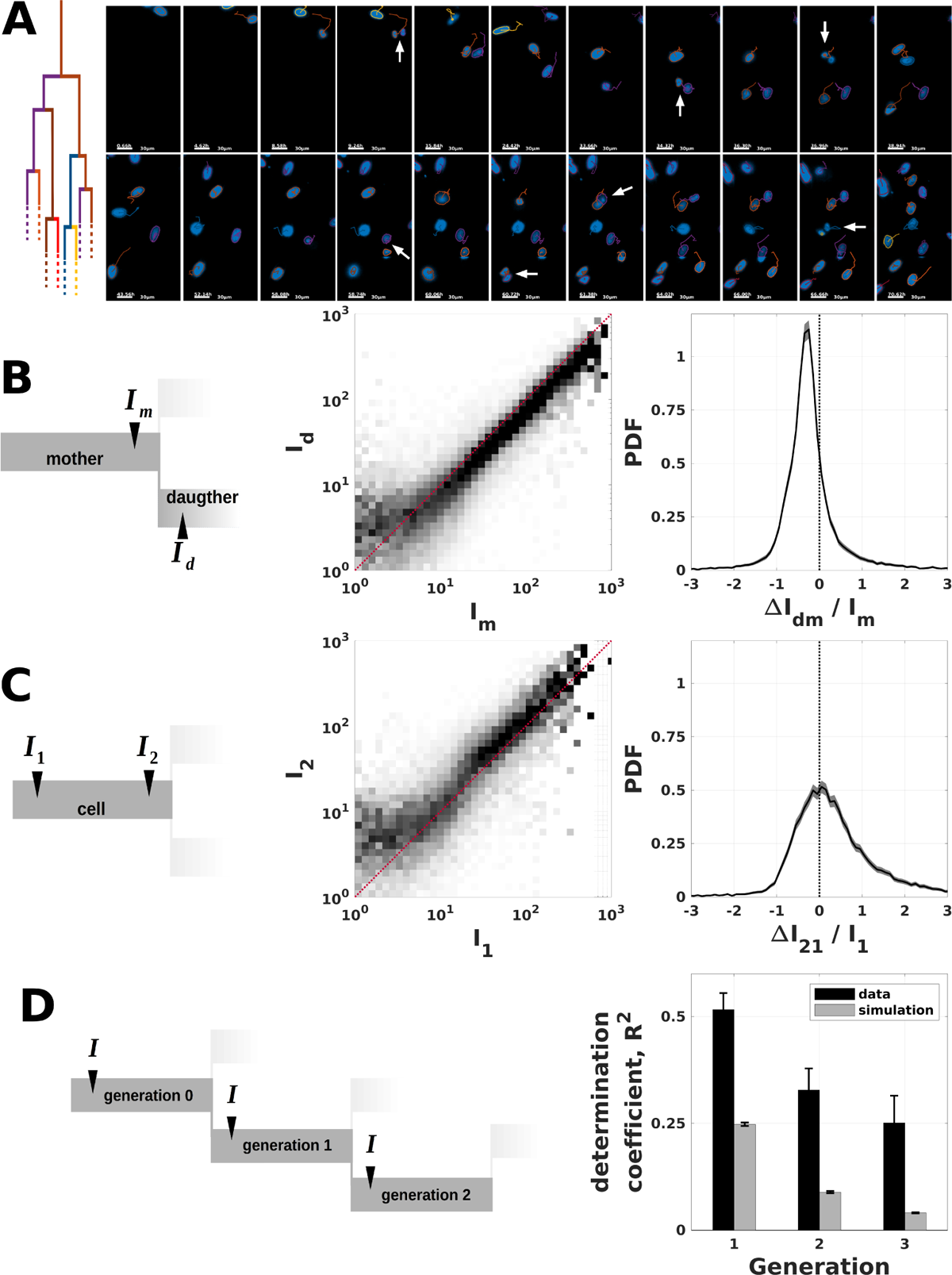
Statistical analysis of phenotypic inheritance of SUM159-PT cells. (**A**) **Representative example of cell lineage:** Hoechst images of a mother cell for which we kept track of its 2 daughters; 4 grand-daughters and 6 out of 8 great grand-daughters. Segmentation masks and cell trajectories are overlaid. Arrows indicate cell division. Scale bar 20*μm*. (**B**) **Correlation of CSC reporter signal between mother and daughters.** Left panel: Mother fluorescence intensity, *I_m_*, is measured at the end of cell cycle and daughter, *I_d_*, fluorescence intensity is measured at the beginning of cell cycle. See materials and methods for details. Middle panel: scatter plot of daugther’s fluorescence as a function of mother’s fluorescence. Right panel: Probability density function of fluorescence variation upon mitosis, Δ*I_dm_* = *I_d_* − *I_m_*, relative to mother’s fluorescence, *I_m_*. Black line represents the estimated PDF. Gray shaded area is the 99% confidence interval obtained by bootstrap resampling (sensitivity analysis, see materials and methods). (**C**) **Correlation of CSC reporter signal between beginning and end of cell cycle for the same cell.** Left panel: Fluorescence intensity is measured both at the beginning of cell cycle (*I*_1_) and at the end (*I*_2_). See materials and methods for details. Middle panel: scatter plot of *I*_2_ as a function of *I*_1_. Right panel: Probability density function of fluorescence variation during cell cycles, Δ*I*_2_ = *I*_2_ − *I*_1_, relative to fluorescence at beginning of cell cycle, *I*_1_. Black line represents the estimated PDF. Gray shaded area is the 99% confidence interval obtained by bootstrap resampling (sensitivity analysis, see materials and methods). (**D**) **Correlation of CSC reporter signal across generations.** Left panel: Fluorescence intensity, *I*, is measured both at the beginning of cell cycle for each cell. See materials and methods for details. Right panel: Determination coefficient between signal of a mother cell and signal of its progeny as a function of generation (1: daugthers; 2: grand-daughters and 3: great grand-daughters). Black bars correspond to data and gray bars correspond to numerical simulation of a memory-less chain model (see materials and methods). Error bars represent the 99% confidence interval obtained by bootstrap resampling (sensitivity analysis, see materials and methods). Figure 4—figure supplement 4. Repeatability of fluorescence variations during cell cycle and upon mitosis in SUM159PT cells Figure 4—figure supplement 4. symmetric and asymmetric division rates of SUM159PT cells Figure 4—figure supplement 4. Statistical analysis of phenotypic inheritance of MDA-MB-231 cells

We first consider fluorescence correlation between mothers and daughters. To do so, we compared the fluorescence intensity measured at the end of the mother’s cycle with the one measured at the beginning of the daughter’s cycle (see materials and methods). From the 5 experiments with SUM159-PT cells, we could retrieve 37,582 of such relationships. As expected daughter’s fluorescence strongly correlates with the one of its mother (Figure 4B center panel). The determination coefficient, which measures the fraction of variance explained by linear correlation, is *R*^2^ = 0.70. Indicating that 70% of the daughter’s signal variance is explained by a linear relationship with mother’s signal. However, fluorescence variation during mitosis is systematically biased toward negative values, indicating that cell mitosis may coincide with chromatin changes that could impact either ALDH1A1 promoter transcription activity or protein degradation (Alber et al., 2018). Relative fluorescence variation indeed shows a bell shape distribution centered around −0.25 (right panel of Figure 4B). This behaviour is reproducible considering each experiment separately (figure Supplement 1A). Repeating the same analysis for MDA-MB-231 cells (figure Supplement 3A) shows a weaker but significant correlation between mother and daughter with *R*^2^ = 0.39. Importantly, divisions are mostly symmetric (see figure Supplement 2). Indeed, fluorescence of two sister cells are strongly correlated (*R*^2^ = 0.66 for SUM159PT cells).

We then revisited the measurements of fluorescence variations during cell cycle by comparing signal measured at the beginning of cell cycle with the one measured at the end (center panel of Figure 4C). We used the 19620 SUM159PT cells for which the full cell cycle is monitored. The determination coefficient is *R*^2^ = 0.59, indicating a weaker correlation between these two signals compared to the correlation between mother and daughter. As opposed to signal variation during mitosis, fluorescence variation during cell cycle is distributed around zero but with a tail biased toward higher values (right panel of Figure 4C). Indeed, ∼ 30% of the cells at least double their fluorescence level. These data are in agreement with the time series clustering from which we found ∼ 30% of cells in transiting clusters. This behaviour is reproducible considering each experiment separately (figure Supplement 1B). Analysis of MDA-MB-231 reveals higher variability of phenotype during cell cycle with *R*^2^ = 0.16 (figure Supplement 3B).

Finally, to better understand how the fluorescent phenotype is transmitted across generations, we computed the determination coefficient between the fluorescence intensities of the mother cell and its progeny of first, second and third generation (Figure 4D). The fluorescence level is measured at the beginning of cell cycle for each generation. As expected, correlation decreases as generation increases with a determination coefficient of *R*^2^ = 0.51 for generation 1, *R*^2^ = 0.32 for generation 2 and *R*^2^ = 0.25 for generation 3. Repeating the same analysis for MDA-MB-231 cells we found similar behavior but with weaker correlation between a cell and its daughters (*R*^2^ = 0.16). Interestingly, the correlation between a cell and its progeny is found to be stronger than in case of a memoryless chain of processes (Figure 4 D). In such a memory-less model, phenotypic inheritance upon division only depends on the state (fluorescence, I_m_) of the mother cell and, similarly, fluorescence variation during cell cycle (I_2_ − I_1_) only depends on fluorescence at the beginning of cell cycle (I_1_) (see material and methods). The observed correlation between a cell and its progeny prompted us to hypothesize a coupling between fluorescence variation during cell cycle and signal decrease after mitosis. We indeed find a weak correlation between ΔI_21_ and ΔI_dm_ (*R*^2^ = 0.35). However, such relationship is fully explained by simulation of a memory-less chain (*R*^2^ = 0.36), ruling out the coupling hypothesis.

Altogether, these data highlight a significant influence of the cell division events on the change of fluorescent signal. Though phenotypic inheritance is observed, the broad distribution of the relative signal increase during the cell cycle reflects the probabilistic nature of differentiating and reprogramming events. In contrast, the relative decrease of signals occurring during mitosis pertains all cells. The strong correlation between a cell and its progeny (that is found to be stronger than in case of a memory-less chain model) suggests the existence of a hidden mechanism that maintain phenotype across cell cycle and upon division.

### Contribution of phenotypic inheritance to the spatial clustering of CSCs

A mechanistic hypothesis to explain the spatial clustering of CSC (Figure 2B-G) relies on the phenotypic inheritance described above (Figure 4D). Divisions where sister cells display similar fluorescence intensities would lead to create spatial correlation due to the proximity of daughter and mother cells. The patterning role of this mechanism depends on the degree of signal correlation between sister cells as well as on the characteristics of cell motility. We need a statistical test to estimate to which extent cell displacement and lineage can solely explain the clustered pattern associated with bivariate PCFs (Figure 2 B-G). To guarantee that the test accurately account for observed cell motility, we used cell trajectories reconstructed from experimental data, while fluorescence variation is obtained from Monte Carlo simulations. At initial time, cells are attributed the fluorescence signal measured from experiment. Time evolution of fluorescence is assumed to be driven by two independent processes (see materials and methods): transitions occur (*i*) upon division (Figure 4B) and (*ii*) during cell cycle (Figure 4C).

On the one hand, simulations reproduce correctly the time evolution of each phenotype at the population level (Figure 5B). After a transient period of ∼ 24 to 36 hours, a steady state is reached with proportions of each phenotype comparable to experimental data. On the other hand, we find a determination coefficient between mother and offsprings of first, second and third generation weaker than the one measured on experimental data (Figure 5C). Similar results are obtained for MDA-MB-231 (figure Supplement 2).

**Figure 5.**
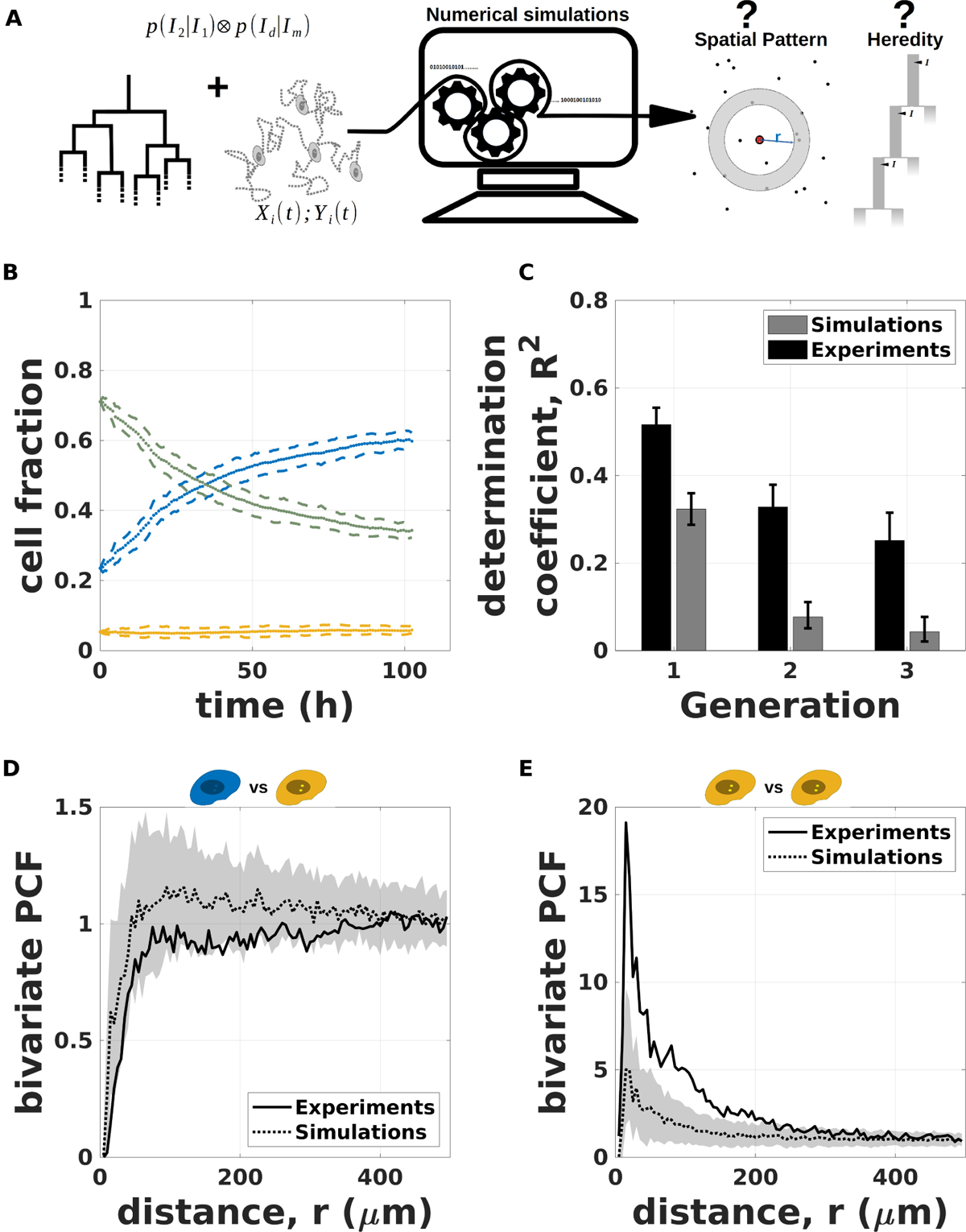
Spatio-temporal simulations of phenotypic inheritance of SUM159PT cells: (**A**) **Simulation workflow:** Cell lineage and cells trajectories are known from experimental data. Probability density function for fluorescence variation are extracted from experimental data. Monte Carlo method is used to simulate fluorescence variation, while cell trajectories are kept identical to experiments. Relevant quantities such as PCF or phenotypic inheritance are computed as for experiments. (**B**) **Simulated time evolution of the cell fraction of the three phenotypes** (CDC blue, iCC green and CSC yellow) defined by the intensity thresholds *I*_−_ and *I*_+_. Mean (points) and 99% confidence interval (dashed lines) obtained from 500 independent simulations based on trajectories of a representative experiment (same as figure Figure 1). (**C**) **Simulated correlation of fluorescence signal across generations.** Is shown determination coefficient between signal of a mother cell and signal of its daughter. Black bars correspond to data and gray bars correspond to numerical simulation (see Material and Methods). Error bars represent the 99% confidence interval obtained by bootstrap resampling (sensitivity analysis, see Materials and Methods). (**D**) **Simulated bivariate PCF for CSC versus CDC.** Continuous black line represents experimental PCF (same as figure Figure 2). Dashed line is the PCF of simulated data for the same experiment. The gray shaded area is the 99% confidence interval obtained by bootstrap resampling (sensitivity analysis, see Material and Methods). (**E**) **Simulated bivariate PCF for CSC versus CSC.** Continuous black line represents experimental PCF (same as figure Figure 2). Dashed line is the PCF of simulated data for the same experiment. The gray shaded area is the 99% confidence interval obtained by bootstrap resampling (sensitivity analysis, see Material and Methods). Figure 5—figure supplement 5. Repeatability of spatio-temporal simulations with SUM159PT cells Figure 5—figure supplement 5. Spatio-temporal simulations of phenotypic inheritance of MDA-MB-231 cells Figure 5—figure supplement 5. Repeatability of spatio-temporal simulations with MDA-MB-231 cells

Importantly, simulations of SUM159PT cells fail to reproduce the spatial clustering of CSC (Figure 5E). For instance, the maximum bivariate PCF, *g*_++_, (found at *r* ≈ 15 μm), is only 3 to 5 times greater than shuffing for all Monte-Carlo simulations, whereas the experimental value is 10 to 25 times higher than shuffing (Figure 5E and figure Supplement 1C). This behaviour is reproducible from one experiment to another (figure Supplement 1C) with p-values below 0.01 for all experiments. Moreover, simulations of SUM159PT cells fail to explain mutual exclusion of CSC and CDC (Figure 5D and figure Supplement 1A) or mutual exclusion of iCC and CDC (figure Supplement 1D). Indeed, bivariate PCF, *g*_+−_(*r* ≈ 15 μm), is only 2 fold lower than shuffing for all Monte-Carlo simulations, while it is more than 10 fold lower for all experiments. Simulations results are more mitigated for MDA-MB-231 cell line for which we found more variability from one experiment to another (figure Supplement 3A-I). We, however, note that simulations fail to reproduce mutual exclusion of CSC and CDC (figure Supplement 3A and I) with p-values below 0.05 for all experiments.

Altogether, these data suggest that phenotypic inheritance of progeny that tends to be colocalized is not enough to explain the observed spatial pattern, probably because of cell motility and the progressive loss of phenotypic memory. Another mechanism is therefore required to promote the formation of CSC clusters, which presumably involves phenotypic transitions driven by environment-specific cues.

### Contribution of intercellular coupling to phenotypic transitions and CSC reprogramming

Motility and division processes are not sufficient to fully explain the spatial patterning of cancer cell phenotypes. Moreover, phenotypic inheritance across several generations is stronger than expected for memory-less chain (Figure 4D). We hypothesize that a spatial control of phenotypic transitions must therefore occur and promote a clustered cell distribution. CSCs display the most pronounced spatial correlation. This pattern, reminiscent of a niche-like effect, is likely to involve cell-to-cell communication.

Figure 6—video 1 A and B qualitatively shows how CSC niche is progressively established from slow fluorescence increase and phenotypic inheritance. We note that fluorescence is stabilized in cells surrounded by CSC (in the middle Figure 6—video 1 B) of while cells in the vicinity of CDCs seem to exhibit lower signal (upper right of Figure 6—video 1 B). More specifically, focusing on two different lineage descending from the same mother cell (Figure 6—video 1 C) indicates that cell fate may strongly depends on environment.

**Figure 6.**
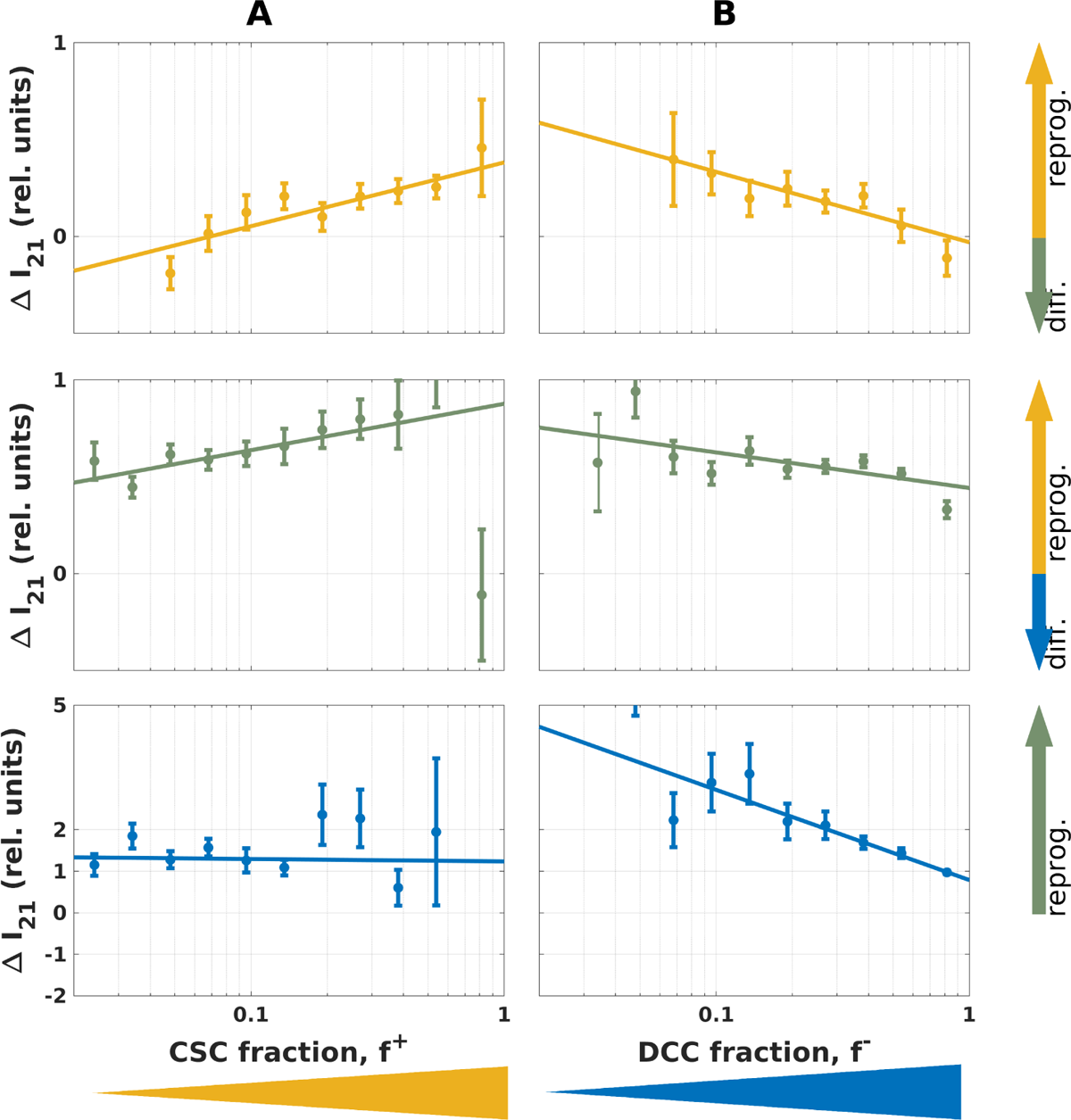
Influence of local environment on fluorescence variation during cell cycle in SUM59PT cells. Average fluorescence variation during cell cycle, Δ*I*_21_ as a function of the local CSC fraction, *f*_+_, (**A**) or the local CDC fraction, *f*_−_, (**B**). From top to bottom, data are shown as function of the initial cell phenotypes: CSC (yellow), iCC (green) and CDC (blue). Δ*I*_21_ is normalized to population average fluorescence intensity of CSC (∼ 150 RFU), iCC (∼ 42 RFU) and CDC (∼ 2.5 RFU). Positive values for Δ*I*_21_ indicate differentiation and negative values indicate reprogramming. Each point represent conditional mean and the height of error bars two standard deviations. Figure 6**—video 1.** Time lapse showing progressive establishment of CSC niche Figure 6—figure supplement 6. Influence of local environment on fluorescence variation upon mitosis in SUM59PT cells Figure 6—figure supplement 6. Influence of local environment on fluorescence variation during cell cycle in MDA-MB-231 cells Figure 6—figure supplement 6. Influence of local environment on fluorescence variation upon mitosis in MDA-MB-231 cells

To investigate the contribution of inter-cellular signaling, we examined fluorescence variation with respect to a quantity reflecting the cell environment. To do so, we computed fluorescence variation conditional to both phenotype and the local spatial density of a given phenotype (see Material and Methods). We considered both the relative density of CSC, f ^+^ and the relative density of CDC, f ^−^.

We first focus on signal variation during cell cycle (ΔI_21_) conditional to spatial environment. Regions of high fraction of CSC phenotypes tend to promote signal increase in iCC and CSC, thus promoting CSC maintenance and iCC reprogramming into CSC (Figure 6A). For instance, for cells classified as CSC at beginning of cell cycle, < ΔI_21_ f ^+^ > is negative (differentiation) when f ^+^ < 0.05 while it increases by 50% (reprogramming) when f ^+^ > 0.9. A consistent trend is obtained in regions with low CDC fraction, except that the signal increase associated to reprogramming extends to CDC phenotypes (Figure 6B, lower panel). For CDC, < ΔI_21_ f ^−^ > value increased by 50% f ^−^ < 0.1 while the value drops close to zero for f ^−^ > 0.9. The MDA-MB-231 cell line also exhibits fluorescence variations that depend on the local cellular environment (figure Supplement 2). In sharp contrast, fluorescence variations upon mitosis (ΔI_dm_) show no significant dependency on the local environment (figure Supplement 1), consistently with a more peaked distribution observed in Figure 4. Similar results are obtained for the MDA-MB-231 cell line (figure Supplement 3). Altogether these results highlight a central role cell-cell communication in spatial patterning and phenotypic inheritance across several generations.

## Discussion

Intra-tumor heterogeneity and cancer stem cells (CSCs) have been recognized to play a role in tumor resistance and invasiveness. Effectively targeting CSC for treatment requires a comprehensive understanding of their multifaceted and intricate plasticity. Despite efforts to characterize the transcriptomic and metabolic profiles of CSCs (REFs), their self-organizing capacity to spontaneously reform stem-like cell niches and spatially-heterogeneous tumors remains poorly understood. In this study, live cell tracking of cultured breast cancer cells marked with a stemness fluorescent reporter enables us to link phenotypic transitions at single-cell level with the emergence of spatiotemporal organization of phenotypic heterogeneity at the population level. ALDH1A expression/activity is commonly used as a CSC marker where high expression is associated to an hybrid Epithelial/mesenchymal CSC phenotype (Grosse-Wilde et al., 2015; Colacino et al., 2018). In our experiments, careful analysis of fluorescent intensity distribution reveals two well-distinct populations characterized with high and low intensities, and therefore classified as CSC-like and CDC-like phenotypes. Interestingly, a population with intermediate level of reporters is also observed and possibly associated with a metastable transiting phenotype or manifesting the existence of diverse non-stem cell types (Colacino et al., 2018).

The phenotype organization suggests a fluid hierarchical arrangement of differentiated states, displaying nevertheless asymmetry between differentiation and dedifferentiation events. Indeed, live cell tracking of fluorescent signal provides insights into the temporal characteristics of both differentiation and dedifferentiation transitions. Temporal resolution of phenotypic transitions is constrained by the time interval of image recording (1 hour) and the degradation timescale of the fused fluorescent proteins (∼ 10h). Given these resolution limitations, dedifferentiation events can be clearly identified during the cell cycle over the course of about ten hours. The timescale of differentiation events are more difficult to extract from signal noise through clustering analysis restricted to one cell cycle. Instead, signs of signal decrease are found systematically just after mitosis which can be due to chromatin condensation and a global transcription arrest during mitosis (Hsiung et al., 2016) or to a symmetric division with accumulation of differentiation factors (Morrison and Kimble, 2006). This suggests that differentiation occurs more progressively at time scales much larger than the cell cycle, or through intermediate or primed states (Sha et al., 2019; Pfeuty et al., 2018). This is overall in good agreement with cell sorting data. Indeed, after selecting only CSC, steady proportions are restored after a long timescale (more than 10 days), while selecting only CDC, new steady proportions are established after only a few divisions (REF thèse justine). Interestingly also, while cancer cell reprogramming into stem-like cells has been previously shown to actively occur in the contexts of radiotherapy or chemotherapy (Lagadec et al., 2012; Auffinger et al., 2014), the present work highlights a significant number of reprogramming events found to occur in an unperturbed cell population. A balance between reprogramming and differentiation events maintains a dynamic equilibrium between heterogeneous phenotypes at the population level.

The advantage of being able to track the phenotypes of a cancer cell population in space and time is to infer the system-level self-organizing mechanism resulting in spatial pattern of cell types. General principles for spatial self-organization relies on diverse cellular processes such as signaling feedbacks, motility, division, where each process involves deterministic and stochastic contributions (Kicheva et al., 2012; Grace and Hütt, 2015; Landge et al., 2020). Spatio-temporal point pattern analysis of cell phenotypes clearly identifies clusters of stem-like cells with a typical size of 150 to 300 μm. The formation of CSC clusters and the spatial exclusion with CDCs are found to originate from the interplay of phenotypic inheritance and signaling cues provided by neighboring cells. All these observations were found in two distinct breast cell lines, SUM159-PT and MDA-MB-231. In both cell lines, symmetric division constitutes a local positive feedback mechanism that is prone to generate clusters of cells with similar phenotypes, while cell-cell interactions constitute population-level feedback that shape and stabilizes long-range spatial structures. In contrast, the contribution of cell motility in cancer cell cultures is not a morphogenetic cell-sorting process (Strandkvist et al., 2014) but rather a source of spatial noise mixing cell phenotypes. Intracellular noise and cell motility thus provide multiple sources of stochasticity influencing the level of spatial correlation and the nature of the patterns. Variation of fluorescent reporter expression and associated noise together with different motile behaviour between cell lines might contribute to the less pronounced spatial segregation and influence of cell to cell interactions found in MDA-MB-231 compared to SUM159-PT.

Can mechanistic signaling insights be inferred from the spatial correlation of phenotypes and phenotypic changesThe clustering of CSC phenotypes is much more marked than for other phenotypes and the reprogramming events are promoted by neighboring CSC cells, which altogether supports a lateral induction mechanism. Lateral induction is a common cell-cell interaction mechanism that can typically generate spatial cell-fate patterns with wavelength of a dozen of cells (Owen et al., 2000; Sjöqvist and Andersson, 2019). In the present case, CSC clustering is observed up to 300μm range. Given a typical diffusivity of *D* ≈ 100 μm^2^*S*^−1^ for a soluble signaling factor. Such a scale is rather compatible with paracrine signaling with a ligand lifetime of 5 to 15*min* (Handly et al., 2015). In contrast, the mutual exclusion between CSC and CDC is observed at a much shorter length scale of 25 to 50 μm which rather supports juxtacrine signaling or paracrine signaling with a ligand of short lifetime or diffusibility. Such a short lifetime is nevertheless unlikely given the low cell density and the geometry of the culture dish in which cells are plated on a surface in contact with a reservoir. The spatial pattern characterized with two very different length scales of the spatial correlations between phenotypes is thus proposed to reflect the involvement of both paracrine and juxtacrine mechanisms in intercellular interactions.

Notch-mediated juxtacrine signaling plays a crucial role in maintaining stemness in cancer stem cells (Meurette and Mehlen, 2018) where Jagged-Notch interactions have been proposed to be involved in the spatial segregation of an hybrid E/M phenotype at the interior of the tumor (Bocci et al., 2019b). Alternatively, self-renewal or differentiation of CSC has been proposed to be sensitive to a wealth of diffusible signaling factors, such as SHH ligands modulating hedgehog pathway (Kim et al., 2013), cytokines modulating JAK/STAT pathway (Ruiz and Altaba, 2011) or DKK1 ligand modulating WNT pathway (Wu et al., 2022).

A mathematical modeling approach would be valuable to test those diverse signaling hypothesis. In particular, naive mathematical models of spatiotemporal phenotypic dynamics in cancer cell population (Olmeda and Ben Amar, 2019) would allow to identify the contribution of diverse intercellular mechanisms to, respectively, the observed spatial patterns and the homeostatic establishment of cell-type fractions. Refining our understanding of the feedback mechanisms that empower cancer stem cells to rapidly reestablish intratumor heterogeneity holds promise for cancer therapy.

## Materials and Methods

### Cell culture, plasmid transfections, and generation of stable cell lines

#### Cell culture

The experiments are carried out on two breast cancer cell lines, SUM159PT and MDA-MB-231. SUM159PT cell line is obtained from Asterand and cultured in F12 Nut Mix media (Gibco) supplemented with fetal bovine serum (5%, FBS, PAN-Biotech), insulin (5 μ*g*∕ml), HEPES (10 nM, 15630080, Gibco), hydrocortisone (1 μ*g*∕ml) and Zell Shield. MDA-MB-231 cell line is obtained from ATCC and cultured in MEM media (Gibco) supplemented with 10% of fetal bovine serum (FBS, PAN-Biotech), 1X of non-essential a*min*o acids (11140035, Gibco) and Zell Shield. Cells were maintained in a 5% CO2/air environment.

#### Plasmid Transfections

The pALDH1A1-mNeptune vector is construct with mNeptune fluorescent protein under control of ALDH1A1 promoter, a breast CSC marker, and is obtained as previously studied (Bidan et al., 2019). Briefly, the high-fidelity DNA polymerase (Q5 DNA polymerase, New England Biolabs) was employed to PCR-amplify the human ALDH1A1 promoter region (−1248 to +52) from genomic DNA of SUM159PT cell line. The mNeptune-TK fused protein coding sequence, replication origin (ori), and neomycin resistance gene were PCR-amplified using various templates vectors. Twenty cycles of PCR were performed, employing specific primers with flanking BsaI sites. The flanking overhangs were selected to complementarily ligate with overhangs from other PCR fragments. Therefore, the PCR fragments can orderly assemble and form a circular vector. The resulting PCR fragments were purified, and close circular plasmids were assembled via a single restriction-ligation reaction with BsaI enzyme (R3535L, New England Biolabs) and T4 DNA ligase (M0202L, New England Biolabs). The assembled plasmids were transformed into competent cells (C404003, Invitrogen). The plasmids were extracted with QIAGEN kits. Sequencing primers were synthesized by Eurogentech and the vector was sequenced by GATC (Sanger sequencing). Thanks to this construction, we can sort the mNeptune high cells that exhibit stemness characteristics (self-renewal, differentiation and tumorigenicity) (Bidan et al., 2019).

The cells were transfected with the vectors using nucleofection. Five hundred thousand cells are resuspended in 100 μ*L* of buffer (kit V, VCA-1003, Lonza) with 1 μ*g* of DNA and electroporated using the X-013 protocol of the Nucleofector II Device (Amaxa).

#### Generation of stable cell lines

Stable cell lines pALDH1A1-mNeptune are obtained as previously explained (Bidan et al., 2019). The mNeptune fluorescence was exa*min*ed 24 hours post-transfection and the positive cells were selected with 1 mg/ml of G418 (Invitrogen). Subsequent to selection, cells positive for fluorescence were sorted using fluorescence-activated cell sorting (BD FACS Aria III). Several days after cell sorting, a heterogeneous cell population with mNeptune positive and mNeptune negative cells is regenerated as expected (Bidan et al., 2019). Regularly, cell cultures are checked for functional stemness reporter. To do so, negative cells are sorted using fluorescence-activated cell sorting (BD FACS Aria III) and we look for establishement of heterogeneous cell population with mNeptune positive and mNeptune negative cells.

After establishment of pALDH1A1-mNeptune stable cell line, cells were transfected with a pCMV-Grx1roGFP2-Hygromycin vector then the stable cell lines were selected with hygromycin B (the vector was modified from pEIGW-Grx1-roGFP2, 64990, Addgene). The information obtained from Grx1-roGFP2 fluorescence is not utilized in this work.

### Time-lapse Microscopy

Ten thousands cells were seeded into 35 mm glass bottom dish in 2 mL of cell media (D35-20-1.5-N, Cellvis). After 24 hours, cells are stained with 1 *μ*g*/ml* of Hoechst 33342 (3570, Invitrogen) in PBS 1X during 20 *min*. Cells were washed with PBS 1X and 3 mL of cell media is added for the time-lapse experiment.

Samples were placed on a Nikon Ti-E microscope equipped with a motorized filters wheel (Nikon) and a XY-motorized stage (Applied Scientific Instrumentation). We used a custom-built top-stage incubator to regulate temperature, humidity, and atmosphere. The incubator was described in (Guilbert et al., 2020). Cells were maintained at 37°C and atmosphere is regulated at 5% *CO*_2_. The microscope and top-stage incubator are placed in an enclosure with temperature maintained at 35°C. This limits temperature gradient between immersion objective and sample.

Cells were imaged on a sCMOS camera (Orca-Flash LT, Hamamatsu) through a 60X microscope objective (NA = 1.4, Nikon). We set the camera binning to 4 resulting in an effective pixel size of 0.43μm. Illu*min*ation for fluorescence and brightfield images was achieved through custom-built optical system (components from Thorlabs) which allows synchronization of illu*min*ation with other apparatus. Exposure time was set to 50 ms for all experiments and for all channels. Hoechst was excited with a 365nm LED (M365LP1, Thorlabs) passing through a band-pass filter (FF01-390/40, semrock) resulting an average light intensity of ∼ 1.0m*W*.mm^−2^ on the sample. Fluorescence was collected and image on the camera *via* a dichroic mirror (FF416-Di01, semrock) and a band-pass filter (FF01-445/20, semrock). m-Neptune was excited with a 590nm LED (M590L3, Thorlabs) passing through a band-pass filter (FF01-593/40, semrock) resulting an average light intensity of ∼ 106m*W*.mm^−2^ on the sample. Fluorescence was collected and image on the camera *via* a dichroic mirror (Di02-R635, semrock) and a band-pass filter (FF01-680/42, semrock).

We use a custom-built acquisition software written in labview to control the setup. The sample is scanned to record 1024 overlapping images (∼222μm × ∼222μm). Channels were recorded sequentially for each given positions. Images stitching lead to a ∼6.8mm × ∼6.8 mm wide field of view. Lateral overlap between high-resolution images was 10μm in both directions. Scanning duration was ∼ 50 *min*. And scanning was repeated to perform time-lapse imaging with time resolution of ∼ 50 *min*.

### Image processing, cell segmentation and tracking

Image processing was performed offline using custom written code in Matlab R2019a. Cell segmentation was performed on images of cell nuclei (Hoechst channel) assu*min*g one nucleus per cell. After image restoration, cell segmentation and tracking were performed simultaneously. Tracking data were indeed used to improve segmentation as proposed in (Chalfoun et al., 2016). The SUM159PT dataset was fully curated by human intervention. Cell lineage and single cell fluorescence signal were subsequently extracted.

#### Image restoration

Image restoration aims at correcting shading in-homogeneity. We took advantage of the very high-throughput of our experiments to extract both background and foreground profiles. Such a method have been discussed previously and our methodology is inspired from both (Kask et al., 2016) and (Peng et al., 2017). Prior to shading correction, camera offset was estimated by recording dark images and was then subtracted. Then We use the following expression for shading correction(Kask et al., 2016):

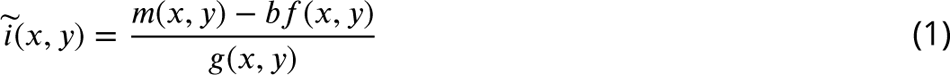

where ĩ and m are respectively an estimation of the true image and the distorted (measured) images, *b* is background intensity, *g* and f are respectively the multiplicative and additive modulating functions.

Thanks to the very high throughput of our experiments, both modulating function, f and *g*, were estimated retrospectively from the data. Indeed, each time-lapse experiment involve recording ∼ 10^5^ vignettes which all have the same modulating functions. In each image, sub-regions could be classified (*via* image segmentation) into either foreground or background. Thus f and *g* could be estimated one after the other by averaging. Modulating function were estimated by combining data from at least 3 different time-lapse experiments. For each experiment, because cells randomly cover the field of view, we separately accumulated data by blocks from either background or foreground. Blocks were averaged to reconstruct both spatial profiles, f and *g*, up to a multiplicative factor (Kask et al., 2016). In order to limit the influence of outliers, we only retain values between 5th and 95th percentiles for both background and foreground. Background was estimated first and non-uniform background was subtracted prior to foreground estimation. For background estimations, images were binned 2 times resulting a 256 × 256 pixels estimation of f. Similarly, for foreground estimation, images were binned 16 times resulting a 32 × 32 pixels estimation of *g*. Averaged modulating functions were then resized to original size (512×512) and smoothed with a Gaussian filter (radius of 2 pixels for background and 16 pixels for foreground). After normalization with respect to their maximum, equation 1 was applied pixel wised. The *b* factor was estimated by averaging m(*x*, *y*)∕f (*x*, *y*) over all background pixels of each image separately.

Masks obtained by segmentation from Hoechst channel were used to reconstruct modulating functions of all channels. Foreground mask was defined as region with identified nuclei while background masks were obtained by excluding disks of radius 22μm around each detected cells. While, given cell segmentation, reconstruction of f and *g* is straightforward, restoration of Hoechst images suffers from a chicken-and-egg problem. To work around this issue, image restoration of Hoechst images was performed in two steps. First, a single modulating function was estimated by assu*min*g the same non-homogeneous profile for background and foreground (f = *g*). A first segmentation was performed which was used only to extract Hoechst channel modulating functions, f and *g*, as described above. Then, the final segmentation of nuclei was performed after restoration using equation 1.

#### Cell segmentation and tracking

We used tracking information to correct segmentation errors as suggested in (Chalfoun et al., 2016). Here, we first describe segmentation then cell tracking and finally explain how tracking is used as a feedback to further refine cell segmentation.

Shading correction was performed as described above. Then, images of nuclei were blurred with a gaussian filter of width 6.5μm to remove tiny details useless for nuclear shape segmentation. This procedure resulted in bell shaped intensity distributions centered on nuclei. We then detected local maxima to assign putative cell centers. Individual masks were initialized to disks of radius 22μm around putative centers. Nuclear masks were refined by iteratively removing border pixels using Otsu thresholding (Otsu et al., 1975). This procedure is followed by several morphological operations. First, holes are filled, then we performed erosion followed by dilatation with kernel of 2 pixels. Each mask was finally automatically screened to detect putative neighbouring cells for which masks were merged. To split joined masks of neighbouring cells, we compared mask boundary to its convex hull. To do so, for all subset of boundaries found inside convex hull, we selected the point closest to mask center and then computed the ratio between (*i*) distance of this point to center and (*ii*) distance of closest section of convex hull to mask center. If at least two points were found with this ratio smaller than 0.75 the mask was split in two parts.

Cell segmentation as described above was performed for each vignette separately. Then the reconstructed field of view was automatically screened for duplicate cell masks caused by image overlap. Overlapping masks found in two different but neighbouring vignettes were identified as duplicate and only the largest mask was kept for further analysis. Information on nuclei area and Hoechst fluorescence intensity were then collected for the whole field of view and at each time point. These data were then used to filter false positives and, in particular, small masks with low fluorescence signal. Detected cells from all vignettes were assembled to assign coordinates (*X*, *Y*) for tracking. Tracking was performed by connecting all cells detected at a given time point to its nearest neighbour at the next time point. We used the algorithm described in (Sbalzarini and Koumoutsakos, 2005) *min*imizing the following cost function: *c* = ∑_ij_ (*X*_i_ − *X*_j_)^2^ + (*Y*_i_ − *Y*_j_)^2^ + α((*W*_i_ −*W*_j_)^2^ +(w_i_ −w_j_)^2^) where summation runs over all paired cells (i, j), *X*, *Y* are cells coordinates and *W*, w are the lengths of major and *min*or axis of the mask. The parameter α was set to 0.12. Association of cost function higher than 30μm were not considered as described in (Sbalzarini and Koumoutsakos, 2005).

As proposed in (Chalfoun et al., 2016), we used tracking as a feedback to enhance cell segmentation. A common error in nuclei segmentation is that two neighbouring cells come in close contact and lead to detection of a single mask for both cells. We call this event “cells collision”. Collision events can easily be identified from tracking data because the trajectory of one cell prematurely ends. Conversely, the mask from a single cell can be correctly detected at one frame but split in the following frame. We named this event “cell over-split”. Over-split events can be confounded with natural cell division. However, during mitosis the cell transiently become brighter in the brightfield channel. To distinguish between mitosis and over-split we thus used a contrast parameter estimated from brightfield images. We computed histogram of all background pixels. The contrast parameter was defined as the fraction of mask pixels out of the 99% confidence interval. Cells with contrast parameter higher than a user defined threshold (typically 0.2) were considered as mitotic ruling out over-split. Screening the tracking data for over-split or collision events allowed us to correct the segmentation. Tracking was then run again. This procedure was repeated 10 times and the number of collision/over-split events was decaying progressively close to zero.

### Single cell fluorescence distribution and fluorescence thresholds

Shading correction for the CSC reporter channel was performed as described above. After shading correction we applied a median filter in a window of 3 × 3 pixels to remove outlier pixels caused by noise amplification in region of low foreground modulating function. Because the CSC reporter is homogeneously distributed across the cell we used fluorescence averaged over the nuclear region as a proxy for single cell fluorescence, I.

For both cell lines, single cell fluorescence distribution was fitted to a compound distribution: A normal distribution for cells with low intensities (mean μ and standard deviation σ) and a gamma distribution for cells with intermediate intensities (shape parameter k and scale parameter k) and another gamma distribution for cells with higher fluorescence level (shape parameter k_+_ and scale parameter k_+_). The probability density function for single cell fluorescence, I, reads:

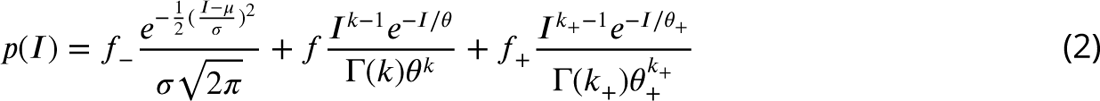

where f_−_, f and f_+_ are respectively the weights of the CDC, iCC and CSC phenotypes. Equation 2 was fitted to data from all experiments all pooled together. We found an optimal set of parameters (all experiments pooled) to be: μ ≈ 2.6, σ ≈ 3.3, k ≈ 2.3, k ≈ 18.2, k_+_ ≈ 2.7 and k_+_ ≈ 55.7 for SUM159-PT. For MDAMB-231 the optimal parameter set was: μ ≈ 47.3, σ ≈ 34.8, k ≈ 3.0, k ≈ 32.4, k_+_ ≈ 1.8 and k_+_ ≈ 115.5. We note that weights of each sub-populations, f_−_, f and f_+_, could vary slightly from one experiment to another.

To attribute phenotype to single cells we used intensity thresholds (I_−_ and I_+_) to separate the three cell populations: cells with signal lower than I_−_ are considered as Differentiated Cancer Cells (CDC); cells with signal higher than I_+_ are considered as Cancer Stem Cells (CSC); and cells with signal between than I_−_ and I_+_ are labelled as intermediate fluorescence Cancer Cells (iCC). Unique pair of thresholds were deter*min*ed based on the fitted distribution for given cell line. I_−_ was chosen so that complementary cumulative distribution function of the CDC cells was bellow 1%. These values were computed independently of the weights, f_−_, and thus depends only on μ and σ. Similarly, I_+_ was chosen so that complementary cumulative distribution function of the CDC cells was bellow 1%. We found I_−_ = 15.5 and I_+_ = 129.5 for SUM159-PT cells and I_−_ = 115 and I_+_ = 245 for MDA-MB-231 cells.

### Data curation

The above segmentation and tracking procedure lead to a false positive rate of *FP* ∼ 5%, a false negative rate of *FN* ∼ 3%, a sensitivity of *S* = *TP* ∕(*TP* + *FN*) ∼ 97% and a positive predictive value of *PPP* = *TP* ∕(*TP* + *FP*) ∼ 95%. While these results are quite good (Caicedo et al., 2019), such an error may be a problem for cell tracking. For instance, assu*min*g an average track length of 20 frames, one expects the probability to detect the cell at all time points to be *S*^20^ = 0.54. In other words, the cell is missed at least once in half of the trajectories. Such a situation will be a problem in particular for lineage reconstruction.

The segmented and tracking data were thus manually corrected using custom-built software in matlab. In brief, segmentation masks were overlaid with images of Hoechst staining and brightfield. Data were screened by a human to detect segmentation errors which were classified into three categories: false positives, missed cells or masks to fuse. Manual correction was saved and corresponding masks were corrected automatically. Tracking was performed once again. Manual correction of a full time-lapse took 40 hours for an untrained user which reduced to 25h after one round. The SUM159PT dataset presented here was fully corrected.

### Lineage reconstruction

Lineage was reconstructed from tracking data by detecting mitotic events. We note that tracking assigns the same identifier to a mother cell and its closest daughter. Putative divisions were detected by screening creation of new trajectory in the vicinity of existing one (distance lower than 60μm). Again we used the contrast parameter estimated from brightfield images to validate mitotic events. If the contrast parameter of either the mother cell or daughter cells was higher than a user defined threshold (typically 0.2) the event was considered as a mitosis.

After detection of all mitotic events, we screened for cells for which we could detect beginning and end of cell cycle, *ie.* cells for which birth and subsequent division were captured by the timelapse and also detected by the algorithm. Only cells with cell cycle duration greater than 15ℎ were retained for time-resolved analysis (lineage analysis and time-series clustering). On the other hand, all cells were used for spatial analysis.

### Point pattern analysis

#### Ripley function and edge correction

The empirical Ripley, K(*r*) function was estimated by counting events within a disk of radius *r* (Cressie, 2015) and averaging over all points of interest. However, observation of the sample in a finite area may lead to biased estimation because information on neighbours for points close to the edge is missing. This effect was corrected by introducing a weight, w_i_(*r*), that rescales counting for points, i, close to the edge (Cressie, 2015). We used uniform correction for which, w_i_(*r*) is the ratio between the area of a disk of radius *r* and the actual observed area within radius *r* (Ripley, 1976, 1977). w_i_(*r*) equals one for points far from the edge and is greater than one for points with truncated observation area.

The empirical Ripley function then reads:

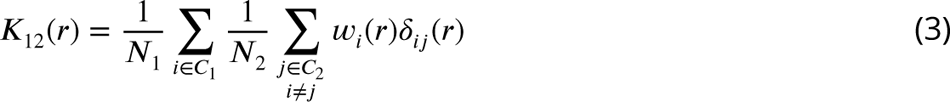

where δ_ij_ (*r*) equals 1 if distance between i and j is smaller than *r* and zero otherwise. C_1_ and C_2_ are two class of points. In the univariate case, C_1_ = C_2_ and N_1_ is the number of points in C_1_ and N_2_ = N_1_ − 1. In the bivariate case, C_1_ ≠ C_2_ and N_1_ and N_2_ are respectively the number of points in C_1_ or C_2_. We note that the Ripley function is symmetrical by construction, *ie.* K_12_ = K_21_.

The Ripley K function was estimated for all *r* up to *r* = 500μm by steps of Δ*r* = 5μm. It was computed in the univariate case (without distinguishing phenotypes) or the bivariate cases where cells were separated in two of the three sub-classes: CSC, iCC, CDC.

#### Point correlation function

The Ripley K function is useful to distinguish clustered or dispersed pattern compared to complete spatial randomness (CSR). However, this function scales with *r*^2^ rendering its visualization and interpretation difficult for all spatial scales together. We instead used the Point Correlation Function (PCF), *g*(*r*) which was estimated as follow (Cressie, 2015):

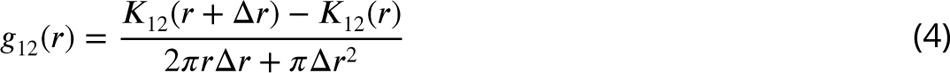

where, K_12_, *r* and Δ*r* were defined above. The PCF, *g*_12_(*r*) can be interpreted as the increase or the decrease of the likelihood of finding an type-2 event at a distance *r* of an type-1 event compared to what would be expected under CSR. *g*_12_(*r*) greater than 1 indicates a more clustered pattern than CSR while a value smaller than 1 indicates a more dispersed pattern than CSR.

### Single cell time-series clustering

The aim of time-series clustering was to identify, without *a priori* knowledge on time scales nor shape, families of single cell temporal patterns of the CSC reporter signal.

To do so, we selected single cell traces for which we could detect beginning and end of cell cycle. Doing so, we obtained 19620 single cell time-series. We used time relative to cell cycle division, τ, to measure cell cycle progression: τ = 0 refers to beginning of cell cycle and τ = 1 to its end. The signals were resampled (zero-order hold resampling (Pohlmann, 2000)) so that all time-series shared the same number of points, N = 60. Euclidean distance, d_kl_, was used to compare single cell time-series: d^2^ = ∑_τ_ (I_k_(τ) − I_l_ (τ))^2^ where I_k_ and I_l_ are signals from two different time series.

All cells drastically change shape during mitosis. They transiently round up and their apparent area was thus smaller than during the rest of cell cycle. This transient morphological change caused a bias in the fluorescence signal estimation during mitosis compared to the rest of the cell cycle. The estimated signal indeed abruptly increased during mitosis and was restored during cycle when the cell was plated back. To avoid clustering on these parts of the time-series, we excluded the six first time points and the six last time points of resampled time-series to search for temporal pattern.

To favor detection of transitioning temporal patterns, clustering was first performed on 6 subsets independently. The 3 first subsets were cells found to be either CDC, iCC or CSC at the beginning of cell cycle and which have changed phenotype at the end; the 3 other subsets were cells that do not change phenotype. We partitioned time series around medoids (Kaufman, 1990; Fränti, 2018). In the initialization step, time series were clustered using a hierarchical procedure where the number of clusters of subset *S*, k_*S*_, was chosen based on a elbow plot. For all subsets, k_*S*_ was between 10 and 25% of the total number of time-series in the subset. A second step aims at optimizing the selection of k_*S*_-centroids. We randomly swapped an existing cluster center (medoid) with a non-medoid time-serie (Fränti, 2018). The permutation was retained if it lead to a decrease of global explained variance, *W*, (Fränti, 2018). This procedure was iterated 1000k_*S*_ times. In a final step, all clusters we merged using a hierarchical procedure.

The final number of clusters, k, was chosen using the Gap-statistics (Tibshirani et al., 2001). This method uses a synthetic datasets to monitor how *W* (k) decreases with k if there where no significant temporal pattern to find. We generated reference datasets of the same size as the experimental one. All time-series of the reference datasets were assumed constant but with additive noise. To do so we randomly initialized synthetic time-series with signal according to the fitted fluorescence distribution. The noise level was chosen by exa*min*ing mean squared deviation of constant experimental traces. The synthetic datasets was then clustered using the same procedure as for the experimental dataset. To select the relevant number of clusters, k^∗^, we compared global explained variance of clustering of experimental data to mean and standard deviation of 20 simulated datasets. As expected, for low values of k, *W* (k) decreases faster for the simulated datasets compared to the experimental data. The optimal number of clusters was chosen as described in the original article (Tibshirani et al., 2001).

### Statistical analysis

#### Bootstrap resampling

Bootstrap resampling (Efron, 1992) was used to estimate sensitivity of several quantities without knowledge of the underlying error distribution. To do so, bootstrap randomly resamples the data with replacement. The procedure is repeated N_*bS*_ times to estimate mean and confidence interval for the quantity of interest. For all uses of bootstrap resampling, we chose N_*bS*_ to ensure that value obtained with bootstrap coincide with the empirical mean estimated without bootstrap(Efron, 1992).

#### Phenotype shuffing

Phenotype shuffing was used as a statistical test for bivariate point pattern analysis. Indeed, the confidence interval of point correlation function (PCF) strongly depends on the number of sample. Moreover, the univariate PCF exhibits a structure indicating that cells display spatial clustering independently of their phenotype.

With phenotype shuffing we aim at deciding whether correlations and anti-correlations between phenotypes revealed by bivariate PCF could be solely explained by the univariate spatial pattern or whether such correlations characterize specific interactions between phenotypes. To do so phenotype shuffing compute phenotype distribution as if phenotypes were randomly distributed across univariate distribution. At a given time point, we used measured cells positions but fluorescence intensities are attributed randomly by permuting all cells intensities. The bivariate PCF are then computed as described above with the simulated intensities. The procedure is repeated 1500 times to compute mean and confidence interval.

#### p-values

p-values are used in this work to assess significance of spatial correlations (anti-correlations) between phenotypes compared to (*i*) phenotype shuffling or (*ii*) numerical simulations. p-values estimate probabilities of the null hypothesis.

For figures 3 and 4, the null hypothesis is “the estimated bivariate PCF does not differ from bivariate PCF upon phenotype shuffling”. To estimate p-values, we first used bootstrap resampling to estimate mean, μ and standard deviation σ of the bivariate PCF at a given radius. Then, we applied phenotype shuffing and computed the corresponding bivariate PCF 1500 times. p-value is defined as the fraction of shuffling for which the corresponding bivariate PCF falls within the range [μ − σ; μ + σ] at the desired radius.

For figures 1 and 3, the null hypothesis is “the experimental bivariate PCF does not differ from the simulated bivariate PCF”. To estimate p-values, we first used bootstrap re-sampling to estimate mean, μ and standard deviation σ of the experimental bivariate PCF at a given radius. We ran spatial simulations and computed the corresponding bivariate PCF 500 times. p-value is defined as the fraction of simulated data for which the corresponding bivariate PCF falls within the range [μ − σ; μ + σ] at the desired radius.

#### Deter*min*ation coefficient

Deter*min*ation coefficient, *R*^2^ is used to exa*min*e correlation between fluorescence signal at different stages of the cell cycle or between a mother cell and its progeny. The deter*min*ation coefficient between random variable *Y* and *X* exa*min* the variance explained by a linear relationship, *Y* ^th^ = a*X* +*b*. Coefficients a and *b* are obtained by fitting the data using matlab built-in fucntion. For a given sample, *R*^2^ reads:

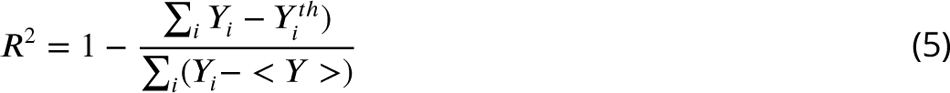

where summation run over all samples and < *Y* > is the sample mean.

### Numerical simulations

Numerical simulations are used for statistical hypothesis testing. Cells are assumed to have an internal variable, I, which represents the fluorescence intensity of the CSC reporter. Cells are assumed to have two states 1 and 2. The first state correspond the beginning of cell cycle and the second to end of cell cycle. Cells can make transition from state 1 to 2 (which correspond to cell cycle evolution) and then from 2 to 1 (which correspond to mitosis).

#### Memory-less chain model

In this model, the internal variable, I, is assumed to change only during state transition. The new value of I depends on its value at the previous state and follows the empirical distribution shown at figure 4 B and C for SUM159PT or figure 3 for MDA-MB-231. To simulate the new value of the intensity, I_2_, at transition 1 ← 2, we first compute the empirical cumulative of the conditional probability density function, *P*(I_2_ I_1_), of I_2_ given I_1_. We used logarithmic sampling for intensity with 62 bins and intensity comprised between −100 and 5000 RFU. A uniform pseudo-random number is generated *via* matlab built-in funciton and I_2_ is obtained by inverse transform of the conditional CDF. The new value of the intensity, I_1_, at transition 2 ← 1, is computed the same but we used the conditional probability density function, *P*(I_d_ I_m_), of I_d_ given I_m_ where I_m_ is replaced by I_1_. Simulations are run 500 times for 10000 cells. Then quantities described in the text are computed the same way as for experimental data.

#### Memory-less spatial model

Motility and divisions are not simulated. Instead we used trajectories and lineage extracted from experiments. Thus for each trajectories, state transition 1 ← 2 and 2 ← 1 are defined by experimentally deter*min*ed lineage. Again, the internal variable, I, is assumed to change only during state transition. Because time evolution of fluorescence of transiting cells was found to be monotonous, we assumed linear time evolution during cell cycle. Evolution of the internal variable, I, is calculated the same way as for Memory-less chain model. Doing so we could simulate intensities at each time point for each cell of the experiment. The simulations were repeated 500 times for each experiments and quantities described in the text are computed the same way as for experimental data.

### Phenotype density estimation

To estimate density of a given phenotype, for each cell, we counted the number of cell within a circle of radius *R* = 300μm centered at the cell position. For cells at the edge, we applied correction as for empirical estimation of the Ripley, K, function. For SUM159PT cells, f_+_ is defined as the number of CSC divided by the total number of cells. Similarly, f_−_ is defined as the number of CDC divided by the total number of cells. For MDA-MB-231 cells, f_+_ is defined as the number of CSC and iCC divided by the total number of cells. This smooth variations given the very low number of CSC in MDA. f_−_ is defined the same way as for SUM159PT.

## Abbreviations

CSCs: Cancer stem cells

iCCs: intermediate cancer cells

CDCs: Cancer differentiated cells

EMT: Epitheial-Mesenchyma Transition

pALDH1A1: promoter of the *ALDH1A1* (Aldehyde Deshydrogenase 1A1) gene

PDF: probability density function

PCF: Point Correlation Function

CSR: complete spatial randomness

SCTS: single-cell time series

GAP: statistics

RFU: Relative Fluorescence Units.

---

**Figure 1—figure supplement 1.**
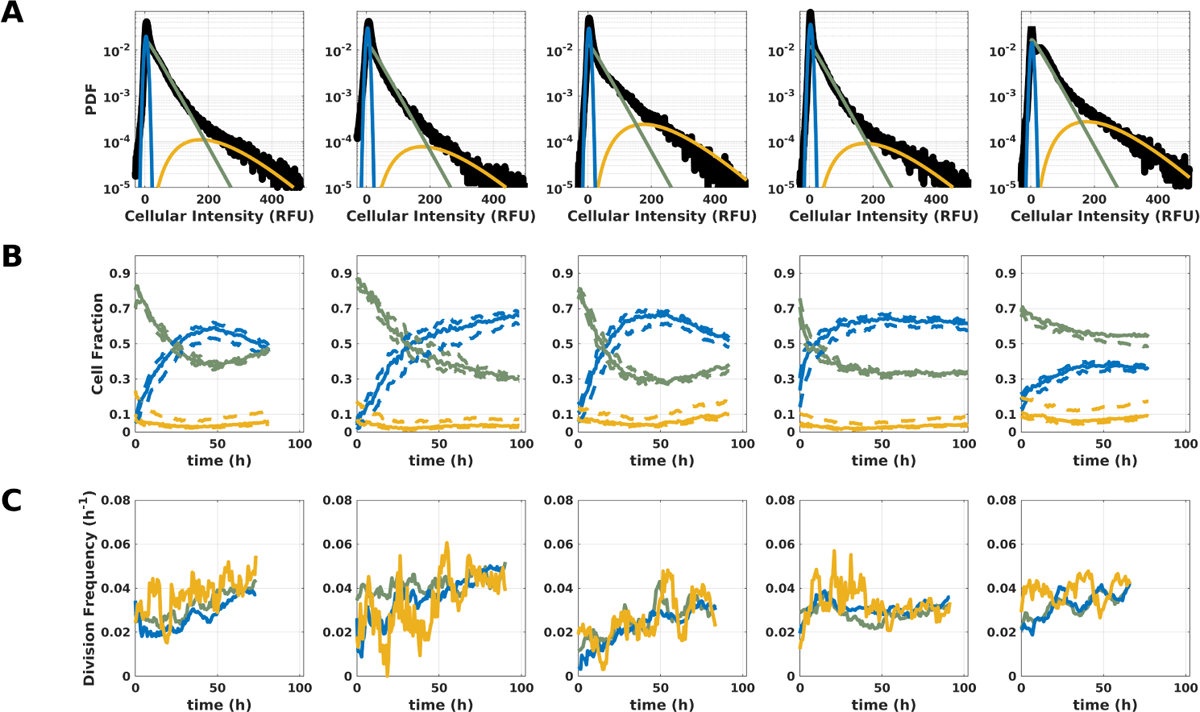
Repeatability of fluorescence distribution and time-evolution in SUM159-PT cells. SUM159-PT breast cancer cells were stably transfected with the CSC reporter, pALDH1a1:mNeptune (Bidan et al., 2019). Cells were imaged as described in the main text. Each column corresponds to an independent experiment. Individual cell nuclei were segmented and the nuclear average signal is used as a proxy for cell phenotype. (**A**) **Fitting of single cell fluorescence distribution.** The probability density function is well fitted by the sum of three distribution (thick black line): A normal distribution for cells with low intensities (blue, mean μ ≈ 2.6 and standard deviation σ ≈ 3.3) and a gamma distribution for cells with intermediate intensities (green, shape parameter k ≈ 2.3 and scale parameter k ≈ 18.2) and another gamma distribution for cells with higher fluorescence level (yellow, shape parameter k_+_ ≈ 2.7 and scale parameter k_+_ ≈ 55.7). Curve fitting was done by Maximum Likelihood Estimation. All experiments were first pooled to deter*min*e shape parameters (μ,σ,k,k,k_+_,k_+_) of each sub-distribution. Then the fitting procedure was repeated for each experiment fixing shape parameters to deter*min*e weights of each distribution.(**B**) **Time evolution of the three populations** (CDCs blue line, iCCs green line and CSCs yellow line) defined by the intensity thresholds (I_−_ = 15.5 and I_+_ = 129.5). Dashed lines show the same but with varying thresholds (12.5≤ I_−_ ≤17.5 and 84.5≤ I_+_ ≤148.5) (**C**) **Instantaneous division rate of all three populations.** We first counted the number of division events detected within a time-windows of 8h for each sub-population. Then, division frequency was estimated by normalizing count by time-window duration and total number of cells of the given sub-population.

**Figure 1—figure supplement 2.**
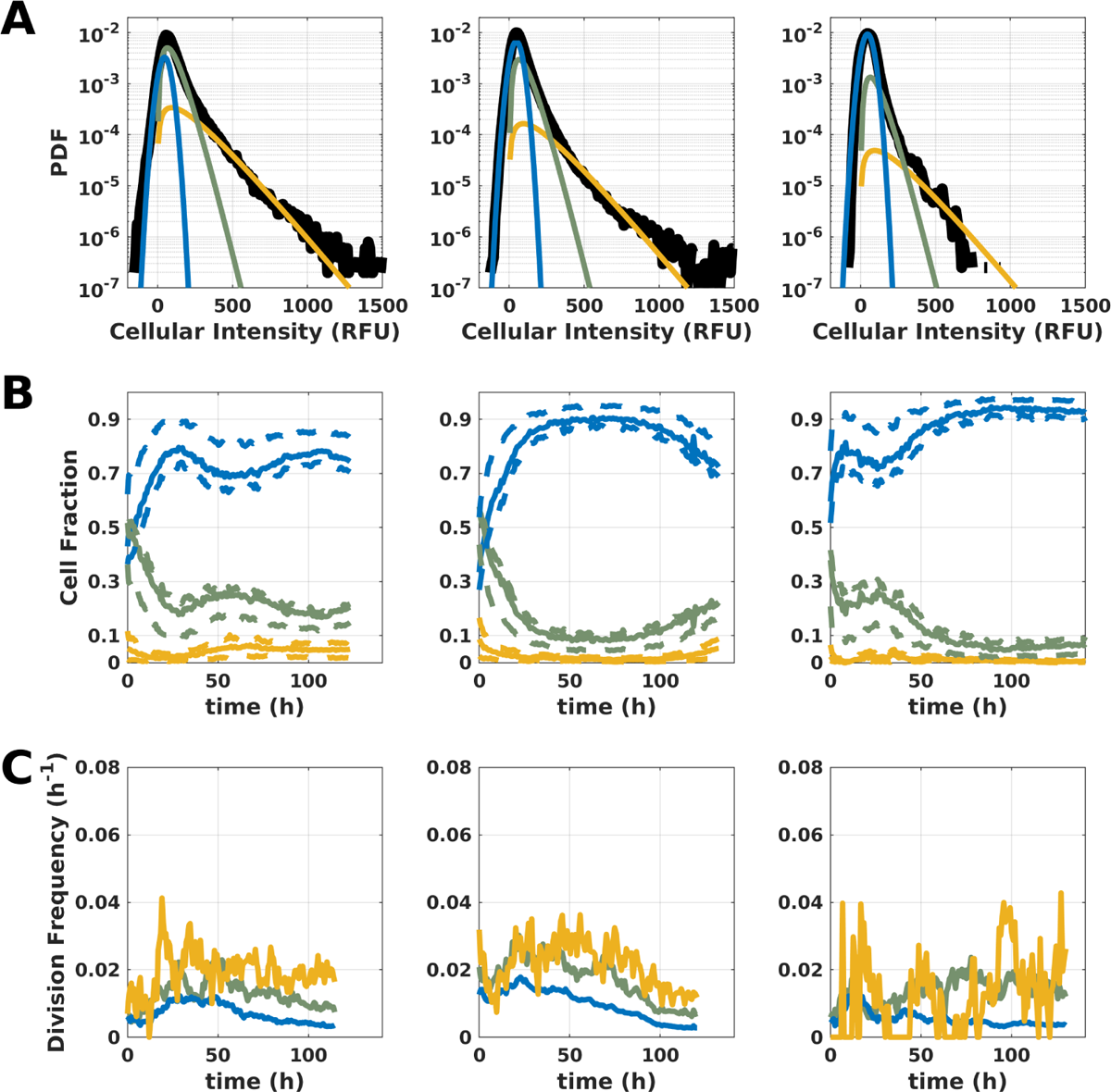
Repeatability of fluorescence distribution and time-evolution in MDA-MB-231 cells. MDA-MB-231 breast cancer cells were stably transfected with the CSC reporter, pALDH1a1:mNeptune (Bidan et al., 2019). Cells were imaged as described in the main text. Each correspond to an independent experiment. Individual cell nuclei were segmented and the nuclear average signal is used as a proxy for cell phenotype. (**A**) **Fitting of single cell fluorescence distribution.** The probability density function is well fitted by the sum of three distribution (thick black line): A normal distribution for cells with low intensities (blue, mean μ ≈ 47.3 and standard deviation σ ≈ 34.8) and a gamma distribution for cells with intermediate intensities (green, shape parameter k ≈ 3.0 and scale parameter k ≈ 32.4) and another gamma distribution for cells with higher fluorescence level (yellow, shape parameter k_+_ ≈ 1.8 and scale parameter k_+_ ≈ 115.5). Curve fitting was done by Maximum Likelihood Maximization. All experiments were first pooled to deter*min*e shape parameters (μ,σ,k,k,k_+_,k_+_) of each sub-distribution. Then the fitting procedure was repeated for each experiment fixing shape parameters to deter*min*e weights of each distribution.(**B**) **Time evolution of the three populations** (CDC blue line, iCC green line and CSC yellow line) defined by the intensity thresholds (I_−_ = 115 and I_+_ = 245). Dashed lines show the same but with varying thresholds (105≤ I_−_ ≤145 and 205≤ I_+_ ≤345) (**C**) **Instantaneous division rate of all three populations.** We first counted the number of division events detected within a time-windows of 8h for each sub-population. Then division frequency was estimated by normalizing count by time-window duration and total number of cells of the given sub-population.

**Figure 2—figure supplement 1.**
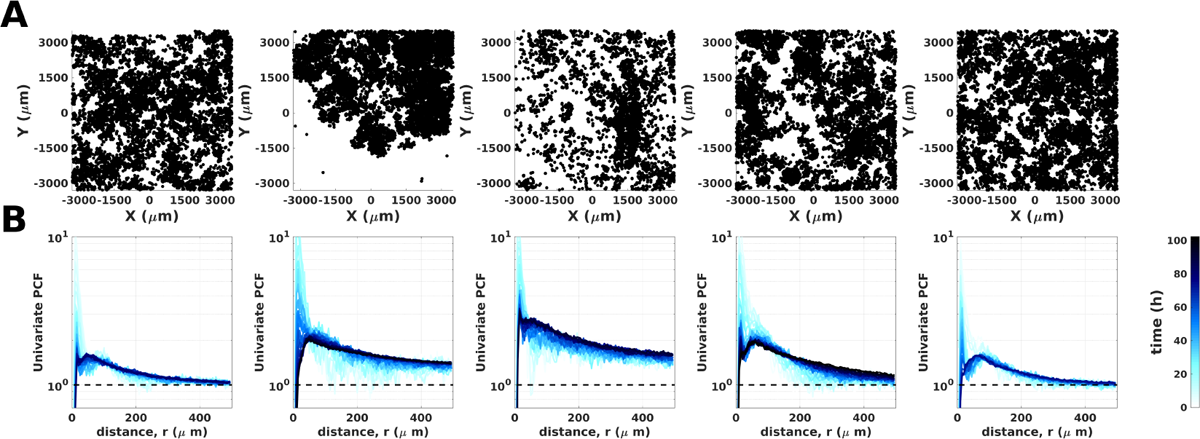
Univariate Point Correlation Function of SUM159-PT cells for all time lapse experiments. (A) Spatial distribution of cells regardless of their phenotype at the end of the experiment. Each cell is represented by a black dot at its spatial coordinates. (B) Univariate PCF for all time points of the time lapse. The PCF, *g*(*r*) is computed at each time point (see Material and methods) and averaged over 5 frames. Line color from light blue to dark blue codes for time (0 to 100h).

**Figure 2—figure supplement 2.**
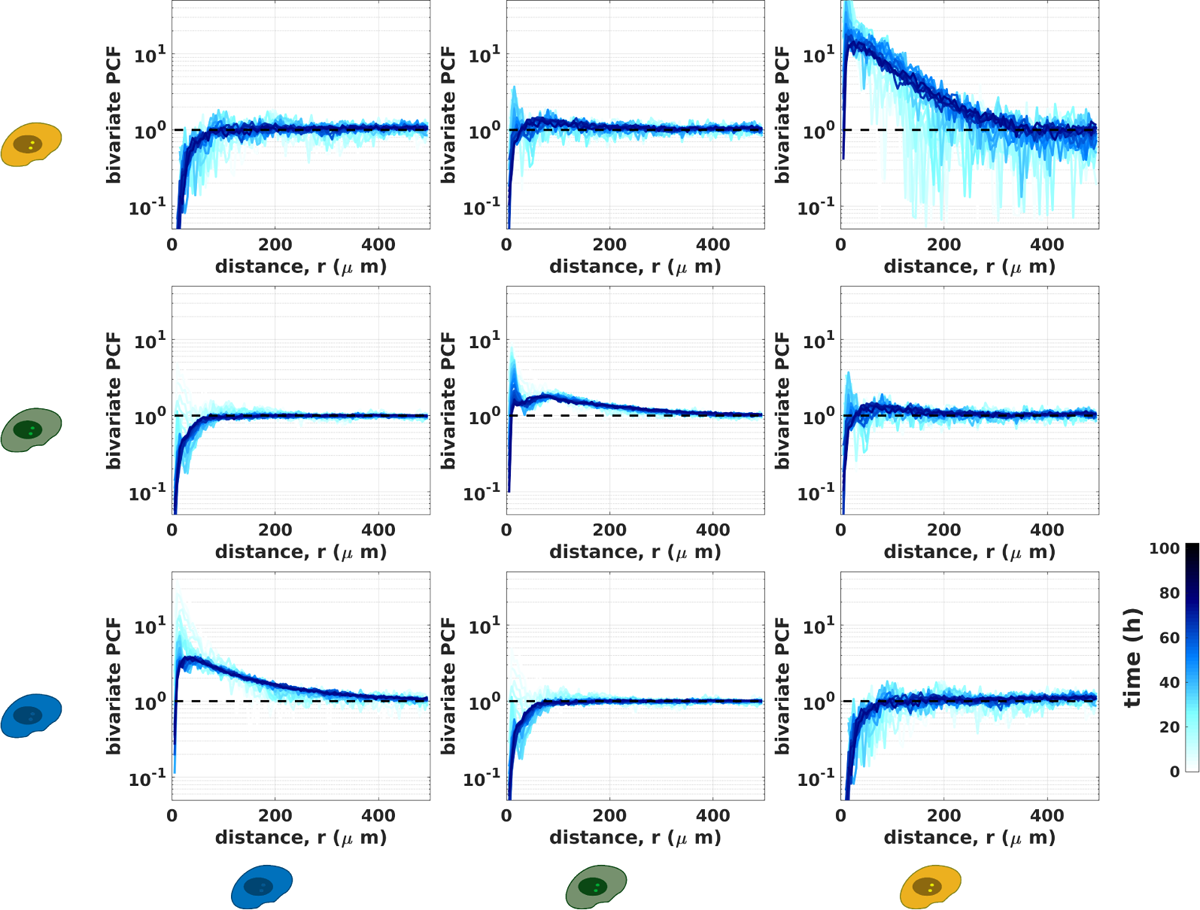
Time evolution of bivariate Point Correlation Functions of SUM159-PT cells for a representative experiment. The experiment is the same as the one shown in main figure. Line color from light blue to dark blue codes for time (0 to 100h). First line: bivariate PCF measuring spatial correlation between CSC and either CDC (first column), iCC (second column) or CSC (third column). Second line: bivariate PCF measuring spatial correlation between iCC and either CDC (first column), iCC (second column) or CSC (third column). Third line: bivariate PCF measuring spatial correlation between CDC and either CDC (first column), iCC (second column) or CSC (third column).

**Figure 2—figure supplement 3.**
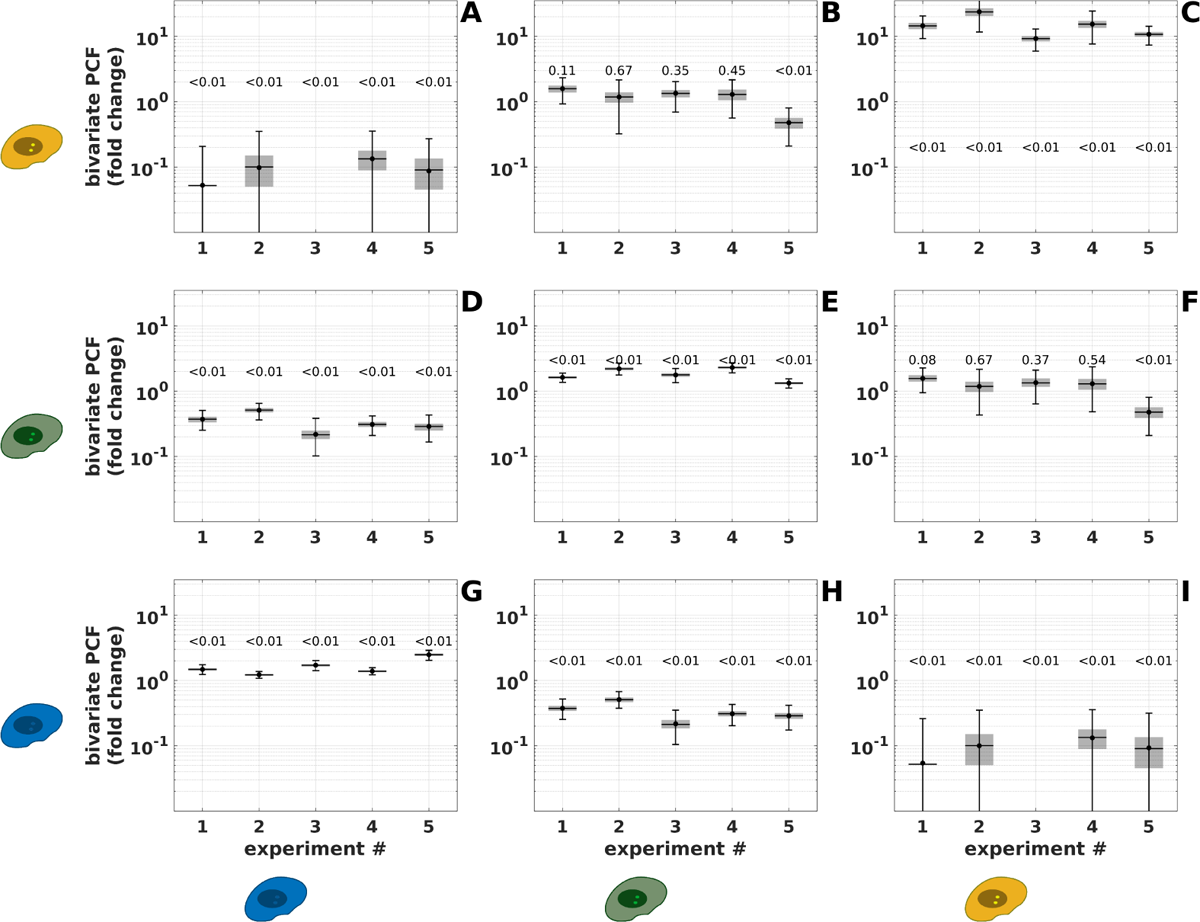
Repeatability of bivariate Point Correlation Function of SUM159-PT cells. We report the ratio between the measured bivariate PCF (data) and the one obtained for phenotype shuffing (control). This ratio is estimated at *r* = 15μm for which the shuffling PCF is maximum. Black dot: actual ratio. Black horizontal line: median obtained from bootstrap resampling. Gray shaded box: 50% confidence interval. Error bars: 99% confidence interval. Note that data point is absent when the measured bivariate PCF is null because of the logarithmic scale. p-values are shown for each data point. p-values are estimated by 1,500 repeats of shuffling to test the null hypothesis (shuffling identical to data within the 68% confidence interval obtained from bootstrap resampling). (**A**) Bivariate PCF measuring spatial correlation between CSC and CDC. Data point is absent for experiment 3 because the measured bivariate PCF is null (cannot be shown on logarithmic scale). (**B**) Bivariate PCF measuring spatial correlation between CSC and iCC. (**C**) Bivariate PCF measuring spatial auto-correlation of CSC. (**D**) Bivariate PCF measuring spatial correlation between iCC and CDC. (**E**) Bivariate PCF measuring spatial auto-correlation between iCC. (**F**) Bivariate PCF measuring spatial correlation between iCC and CSC. (**G**) Bivariate PCF measuring spatial auto-correlation of CDC. (**H**) Bivariate PCF measuring spatial correlation between CDC and iCC. (**I**) Bivariate PCF measuring spatial correlation between CDC and CSC. Data point is absent for experiment 3 because the measured bivariate PCF is null (cannot be shown on logarithmic scale).

**Figure 2—figure supplement 4.**
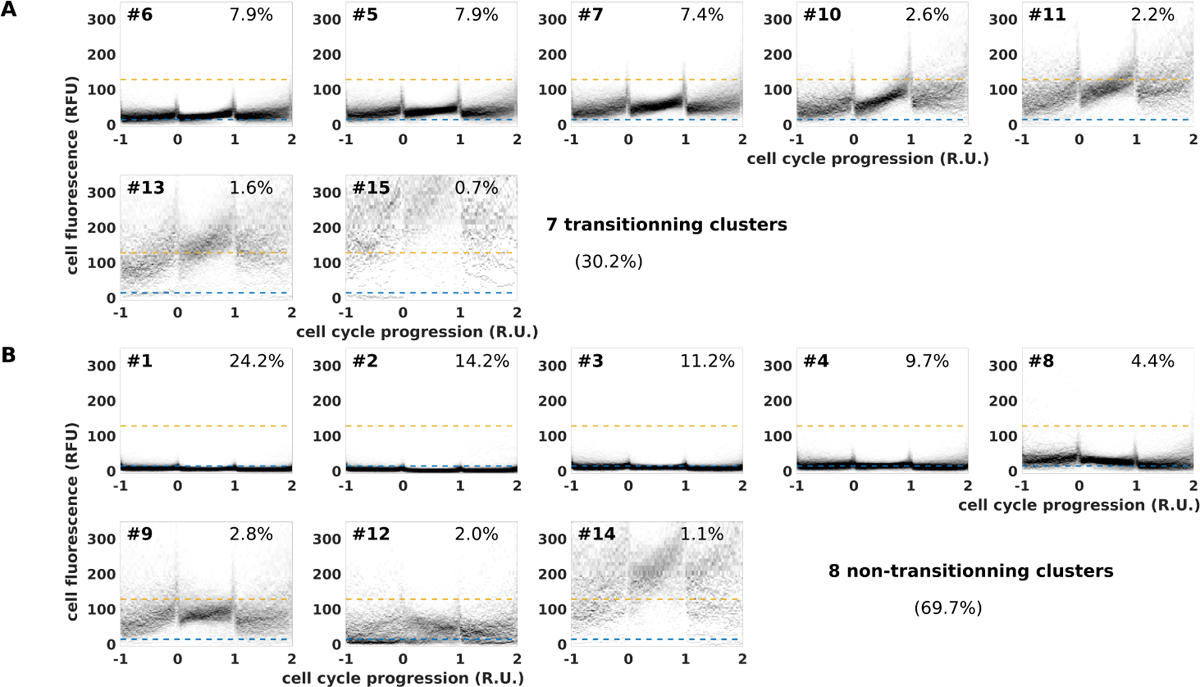
Repeatability of bivariate Point Correlation Function of MDA-MB-231 cells. We report the ratio between the measured bivariate PCF (data) and the one obtained for phenotype shuffing (control). This ratio is estimated at *r* = 15μm for which the shuffling PCF is maximum. Black dot: actual ratio. Black horizontal line: median obtained from bootstrap resampling. Gray shaded box: 50% confidence interval. Error bars: 99% confidence interval. p-values are shown for each data point. p-values are estimated by 1500 repeats of shuffling to test the null hypothesis (shuffling identical to data within the 68% confidence interval obtained from bootstrap resampling). (A) Bivariate PCF measuring spatial correlation between CSC and CDC. (B) Bivariate PCF measuring spatial correlation between CSC and iCC. (C) Bivariate PCF measuring spatial auto-correlation of CSC. (D) Bivariate PCF measuring spatial correlation between iCC and CDC. (E) Bivariate PCF measuring spatial auto-correlation between iCC. (F) Bivariate PCF measuring spatial correlation between iCC and CSC. (G) Bivariate PCF measuring spatial auto-correlation of CDC. (H) Bivariate PCF measuring spatial correlation between CDC and iCC. (I) Bivariate PCF measuring spatial correlation between CDC and CSC.

**Figure 3—figure supplement 1.**
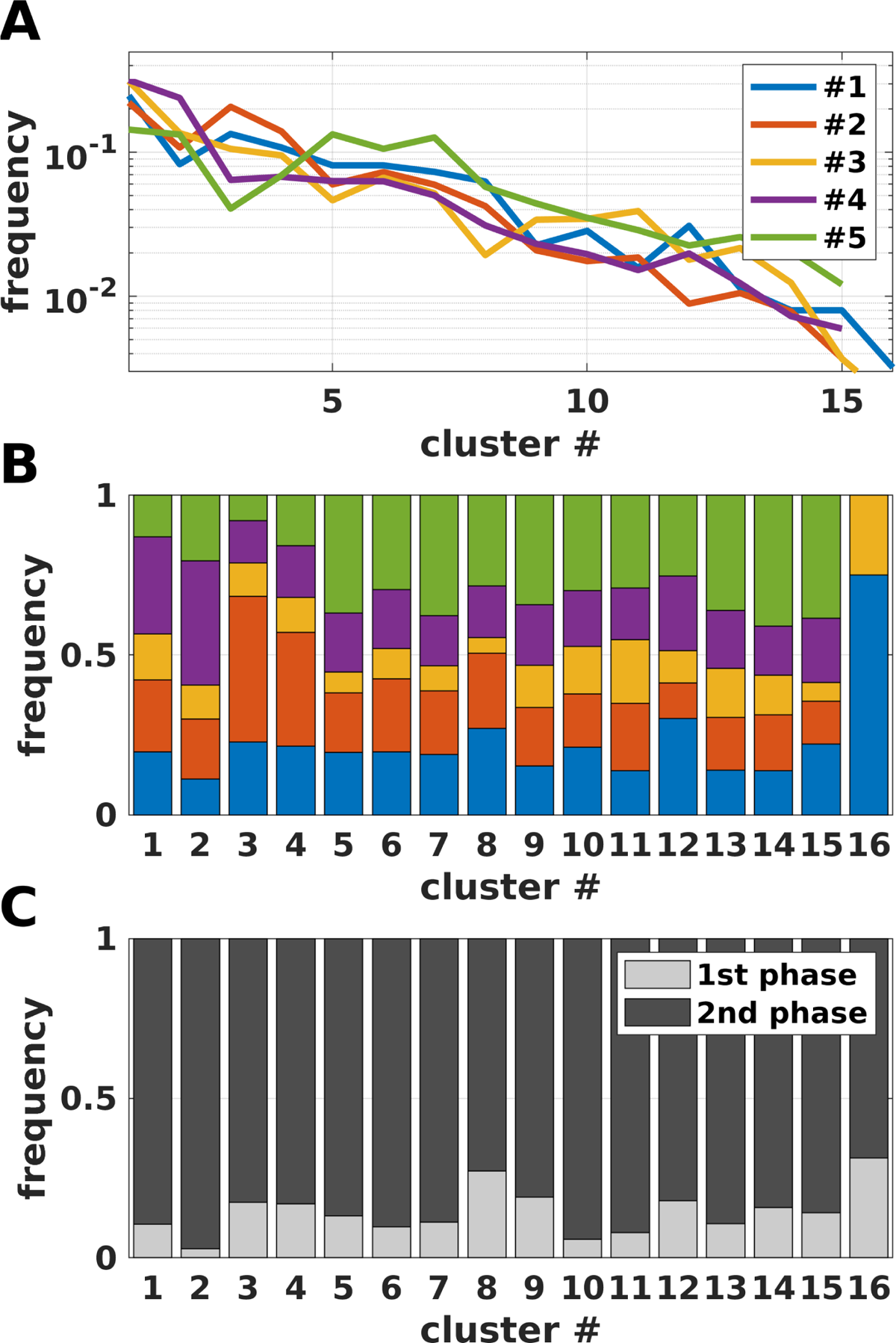
Meta-analysis of identified SCTS clusters: (**A**) Fraction (ordinate) of cells from a given experiment partitioned into a given cluster (abscissa). Each experiment is plotted in a different color. (**B**) Distribution of clusters among each experiment. Color code for experiment number is the same as for A. (**C**) Distribution of clusters among phases of the experiment. First phase is the transient phase and second phase is the stationary one for which steady proportion are reached for all phenotypes.

**Figure 3—figure supplement 2.**
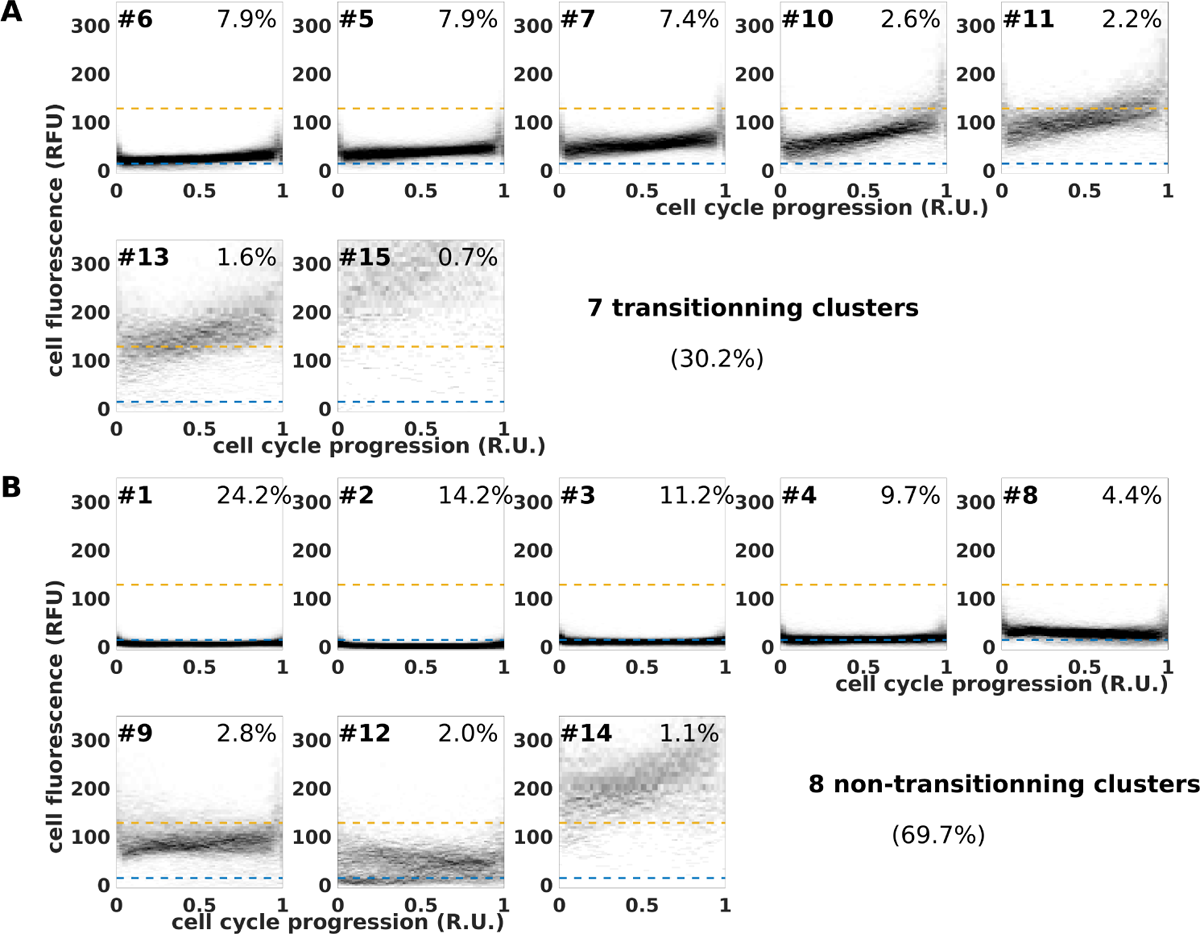
Transitionning (**A**) and non-transitionning SCTS clusters (**B**) in SUM159PT cells. Data are the same as for the main figure. Clusters are numbered by increasing size. The percentage of cells within each cluster is indicated.

**Figure 3—figure supplement 3.**
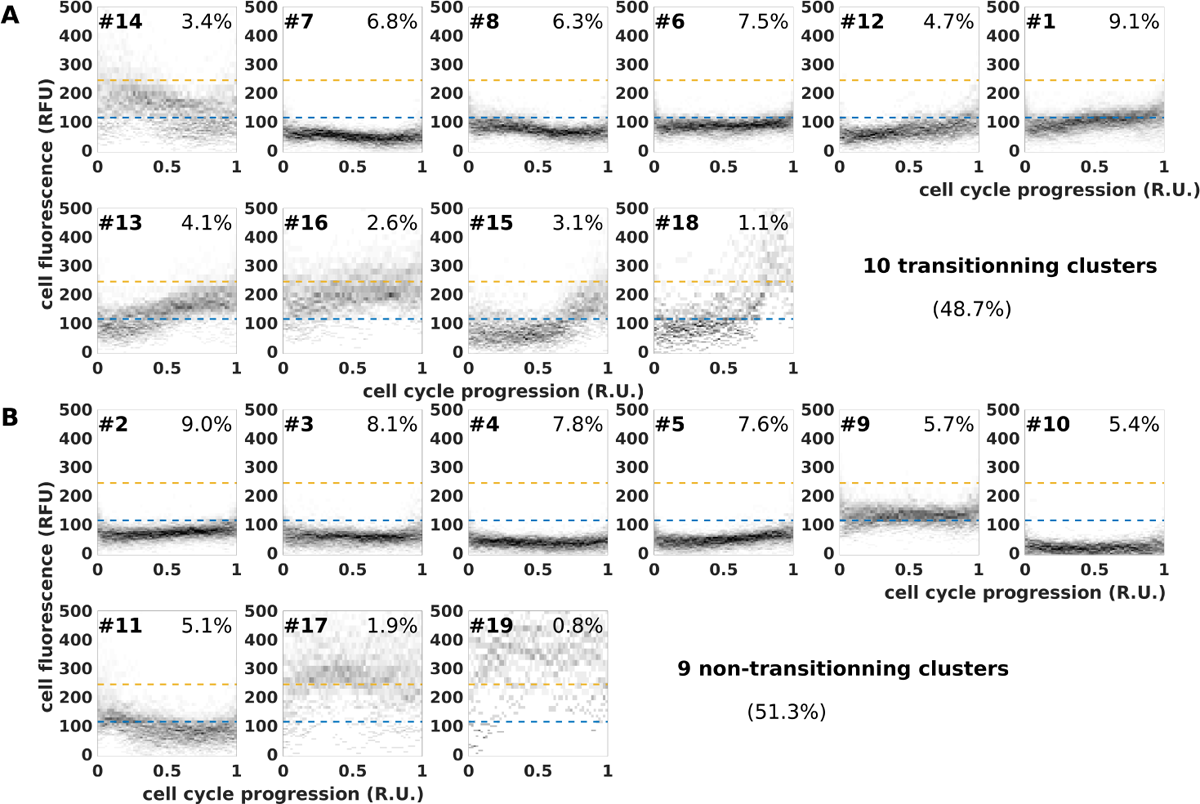
Transitionning (A) and non-transitionning (B) SCTS clusters in MDA-MB-231 cells. Clusters are numbered by increasing size. The percentage of cells within each cluster is indicated.

**Figure 3—figure supplement 4.**
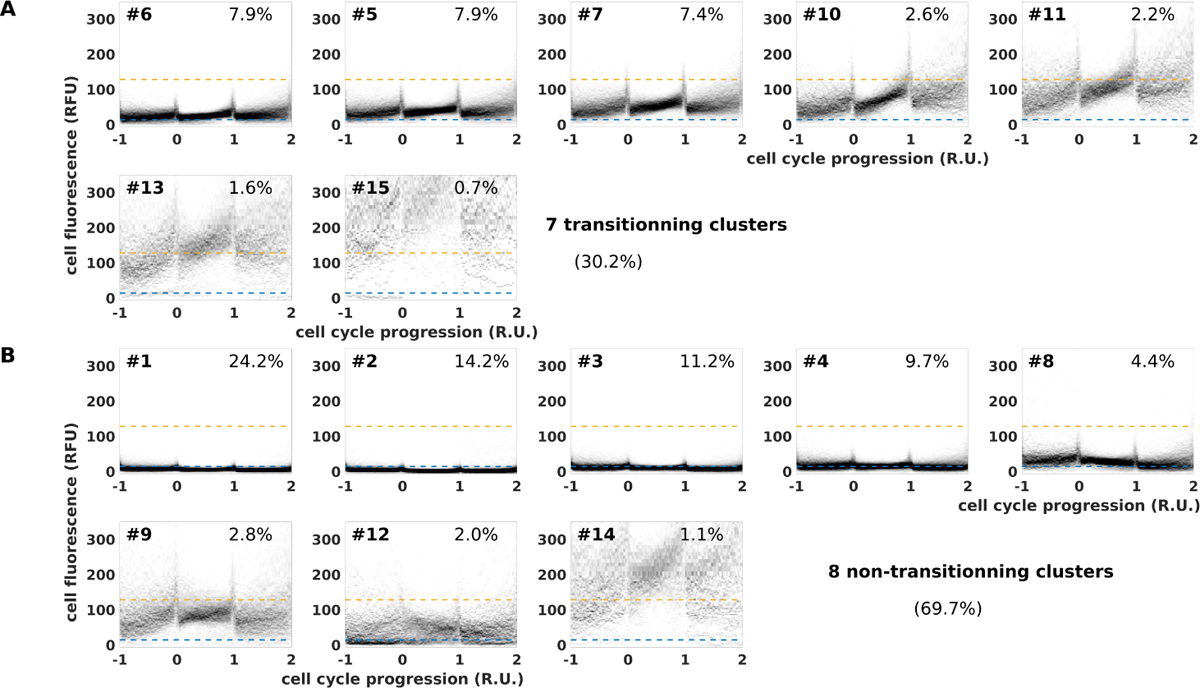
Data are the same as for the main figure but mother signal and daugthers signals are shown when available. Transitionning (**A**) and non-transitionning SCTS clusters (**B**) in SUM159PT cells.

**Figure 4—figure supplement 1.**
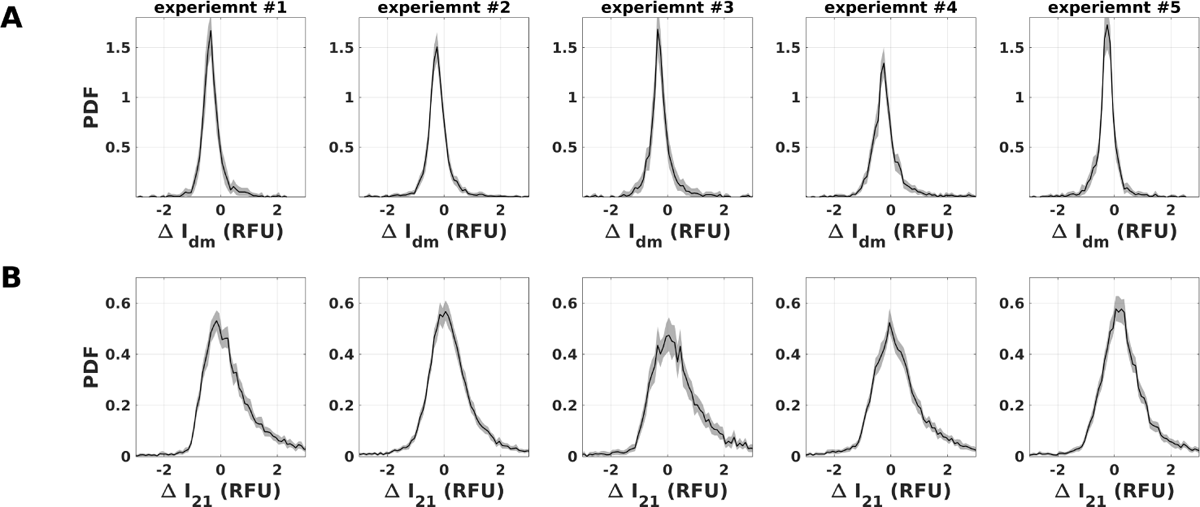
Repeatability of fluorescence variations during cell cycle and upon mitosis in SUM159PT cells: Each column correspond to an experiment.(A) Probability density function of fluorescence variation upon mitosis, ΔI_dm_ = I_d_ − I_m_, relative to mother’s fluorescence, I_m_. Black line represents the estimated PDF. Gray shaded area is the 99% confidence interval obtained by bootstrap resampling (sensitivity analysis, see materials and methods). (B) Probability density function of fluorescence variation during cell cycles, ΔI_2_ = I_2_ − I_1_, relative to fluorescence at beginning of cell cycle, I_1_. Black line represents the estimated PDF. Gray shaded area is the 99% confidence interval obtained by bootstrap resampling (sensitivity analysis, see materials and methods).

**Figure 4—figure supplement 2.**
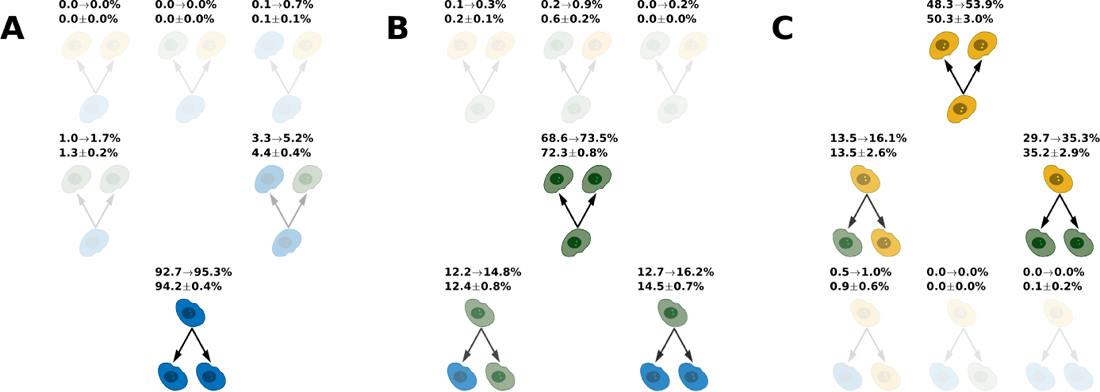
Symmetric and asymmetric division rates of SUM159PT cells: Phenotype are defined according to fluorescence thresholds, I_+_ and I_−_. Blue cell pictogram represents CDC, Green cell pictogram represents iCC and Yellow cell pictogram represents CSC. Percentages indicate rate of a given division type for (**A**) CDC, (**B**) iCC and (**C**) CSC. The lower values represent mean and standard deviation obtained by bootstrap resampling (sensitivity analysis, see materials and methods). The upper interval indicates sensitivity to thresholds: lower and higher values by varying thresholds (12.5≤ I_−_ ≤17.5 and 84.5≤ I_+_ ≤148.5).

**Figure 4—figure supplement 3.**
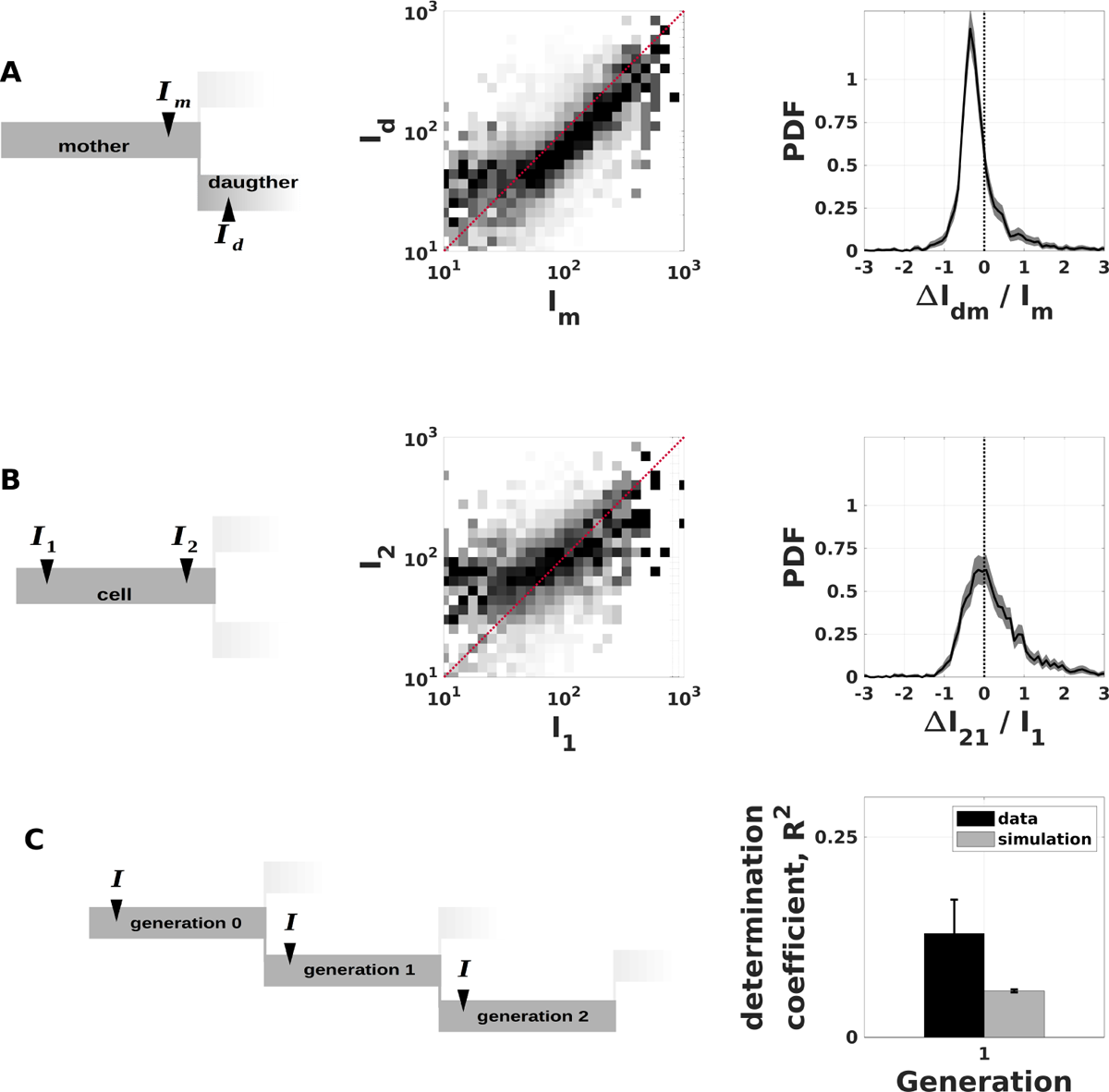
Analysis of fluorescence correlations in the lineage tree of MDA-MB-231 cells. (A) Correlation of CSC reporter signal between mother and daughters. Left panel: Mother fluorescence intensity, I_m_, is measured at the end of cell cycle and daughter, I_d_, fluorescence intensity is measured at the beginning of cell cycle. See materials and methods for details. Middle panel: scatter plot of daughter’s fluorescence as a function of mother’s fluorescence. Right panel: Probability density function of fluorescence variation upon mitosis, ΔI_dm_ = I_d_ − I_m_, relative to mother’s fluorescence, I_m_. Black line represents the estimated PDF. Gray shaded area is the 99% confidence interval obtained by bootstrap resampling (sensitivity analysis, see materials and methods). (B) Correlation of CSC reporter signal between beginning and end of cell cycle for the same cell. Left panel: Fluorescence intensity is measured both at the beginning of cell cycle (I_1_) and at the end (I_2_). See materials and methods for details. Middle panel: scatter plot of I_2_ as a function of I_1_. Left panel: Probability density function of fluorescence variation during cell cycles, ΔI_2_ = I_2_ − I_1_, relative to fluorescence at beginning of cell cycle, I_1_. Black line represents the estimated PDF. Gray shaded area is the 99% confidence interval obtained by bootstrap resampling (sensitivity analysis, see materials and methods). (C) Correlation of CSC reporter signal across generations. Left panel: Fluorescence intensity, I, is measured both at the beginning of cell cycle for each cell. See materials and methods for details. Right panel: Deter*min*ation coefficient between signal of a mother cell and signal of its daughter. Black bars correspond to data and gray bars correspond to numerical simulation of a memory-less chain model (see materials and methods). Error bars represent the 99% confidence interval obtained by bootstrap resampling (sensitivity analysis, see materials and methods).

**Figure 5—figure supplement 1.**
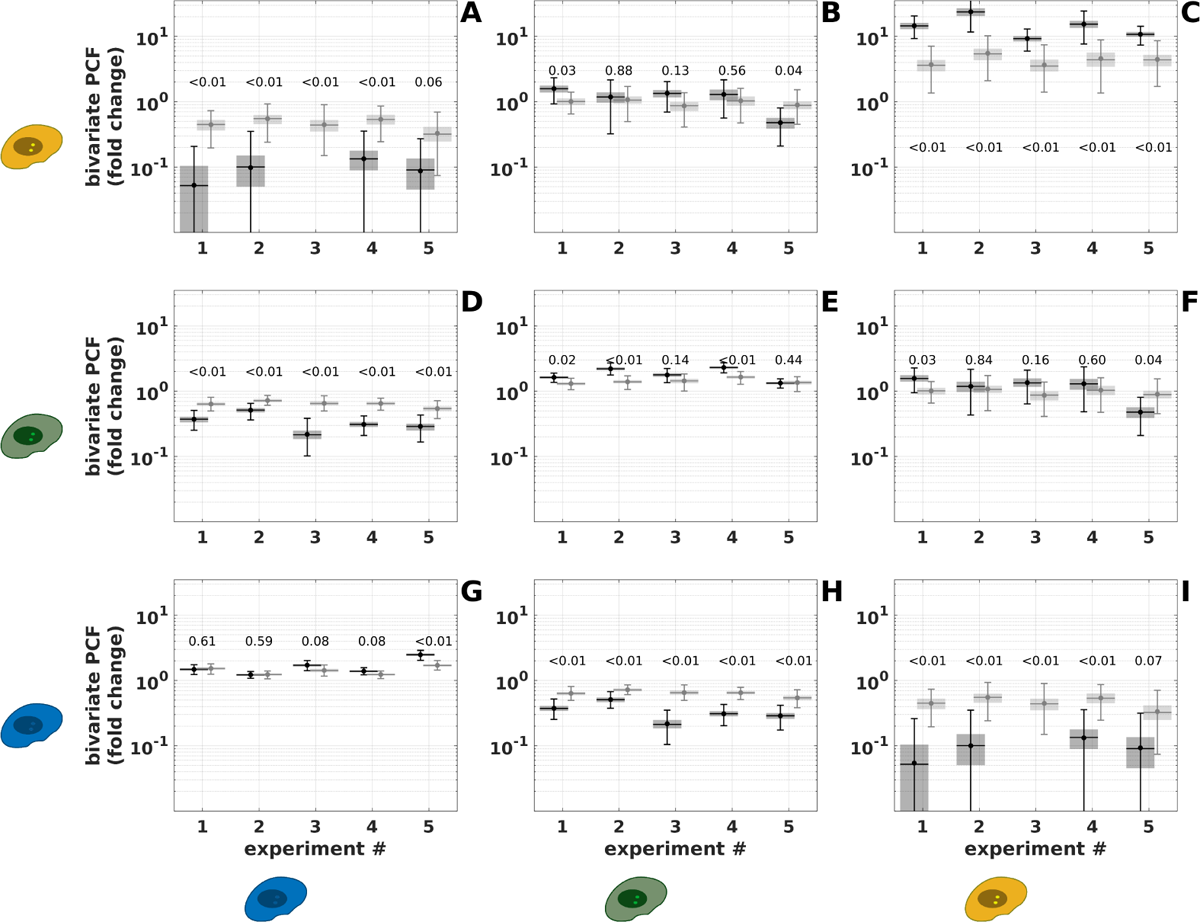
Repeatability of simulated bivariate Point Correlation Function of SUM159-PT cells. We report the ratio between the measured bivariate PCF and the one obtained for phenotype shuffing (control) for data (black) and simulations (gray). This ratio is estimated at *r* = 15μm for which the shuffling PCF is maximum. Dot: actual ratio. Horizontal line: median obtained from bootstrap resampling. Gray shaded box: 50% confidence interval. Error bars: 99% confidence interval. p-values are shown for each data point. p-values are estimated by 500 repeats of simulations to test the null hypothesis (simulations identical to data within the 68% confidence interval obtained from bootstrap resampling). (**A**) Bivariate PCF measuring spatial correlation between CSC and CDC. Data point is absent for experiment 3 because the measured bivariate PCF is null (cannot be shown on logarithmic scale). (**B**) Bivariate PCF measuring spatial correlation between CSC and iCC. (**C**) Bivariate PCF measuring spatial auto-correlation of CSC. (**D**) Bivariate PCF measuring spatial correlation between iCC and CDC. (**E**) Bivariate PCF measuring spatial auto-correlation between iCC. (**F**) Bivariate PCF measuring spatial correlation between iCC and CSC. (**G**) Bivariate PCF measuring spatial auto-correlation of CDC. (**H**) Bivariate PCF measuring spatial correlation between CDC and iCC. (**I**) Bivariate PCF measuring spatial correlation between CDC and CSC. Data point is absent for experiment 3 because the measured bivariate PCF is null (cannot be shown on logarithmic scale).

**Figure 5—figure supplement 2.**
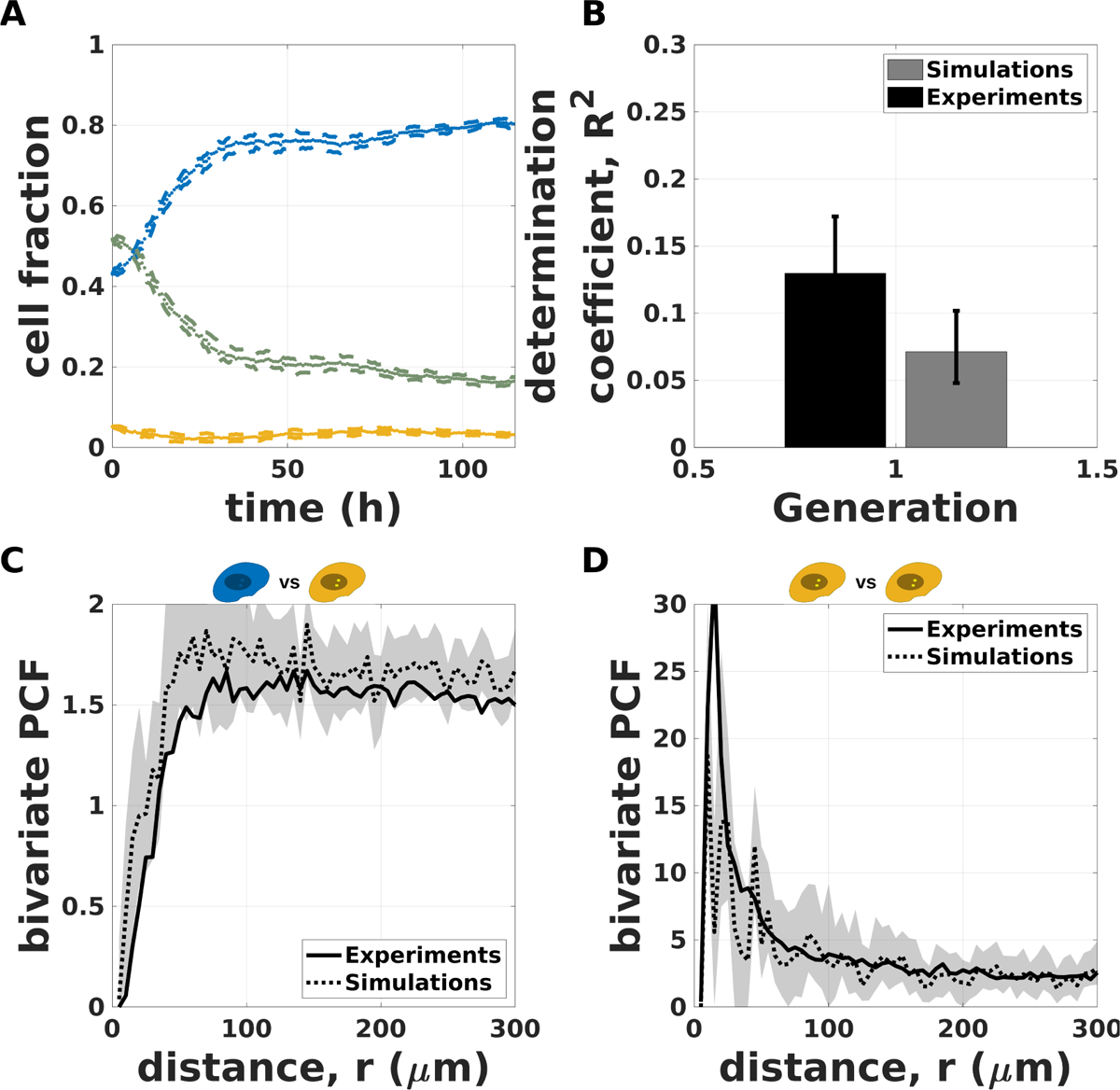
Spatio-temporal simulations of phenotypic inheritance of MDA-MB-231 cells (A) Simulated time evolution of the cell fraction of the three phenotypes (CDC blue, iCC green and CSC yellow) defined by the intensity thresholds I_−_ and I_+_. Mean (points) and 99% confidence interval (dashed lines) obtained from 500 independent simulations based on trajectories of a representative experiment (same as figure *Figure 1*—*figure Supplement 2*). (B) Simulated correlation of fluorescence signal across generations. Is shown deter*min*ation coefficient between signal of a mother cell and signal of its daughter. Black bars correspond to data and gray bars correspond to numerical simulation (see material and methods). Error bars represent the 99% confidence interval obtained by bootstrap resampling (sensitivity analysis, see material and methods). (C) Simulated bivariate PCF for CSC versus CDC. Continuous black line represents experimental PCF (same as figure *Figure 2*—*figure Supplement 4*). Dashed line is the PCF of simulated data for the same experiment. The gray shaded area is the 99% confidence interval obtained by bootstrap resampling (sensitivity analysis, see material and methods). (D) Simulated bivariate PCF for CSC versus CSC. Continuous black line represents experimental PCF (same as figure *Figure 2*—*figure Supplement 4*). Dashed line is the PCF of simulated data for the same experiment. The gray shaded area is the 99% confidence interval obtained by bootstrap resampling (sensitivity analysis, see material and methods).

**Figure 5—figure supplement 3.**
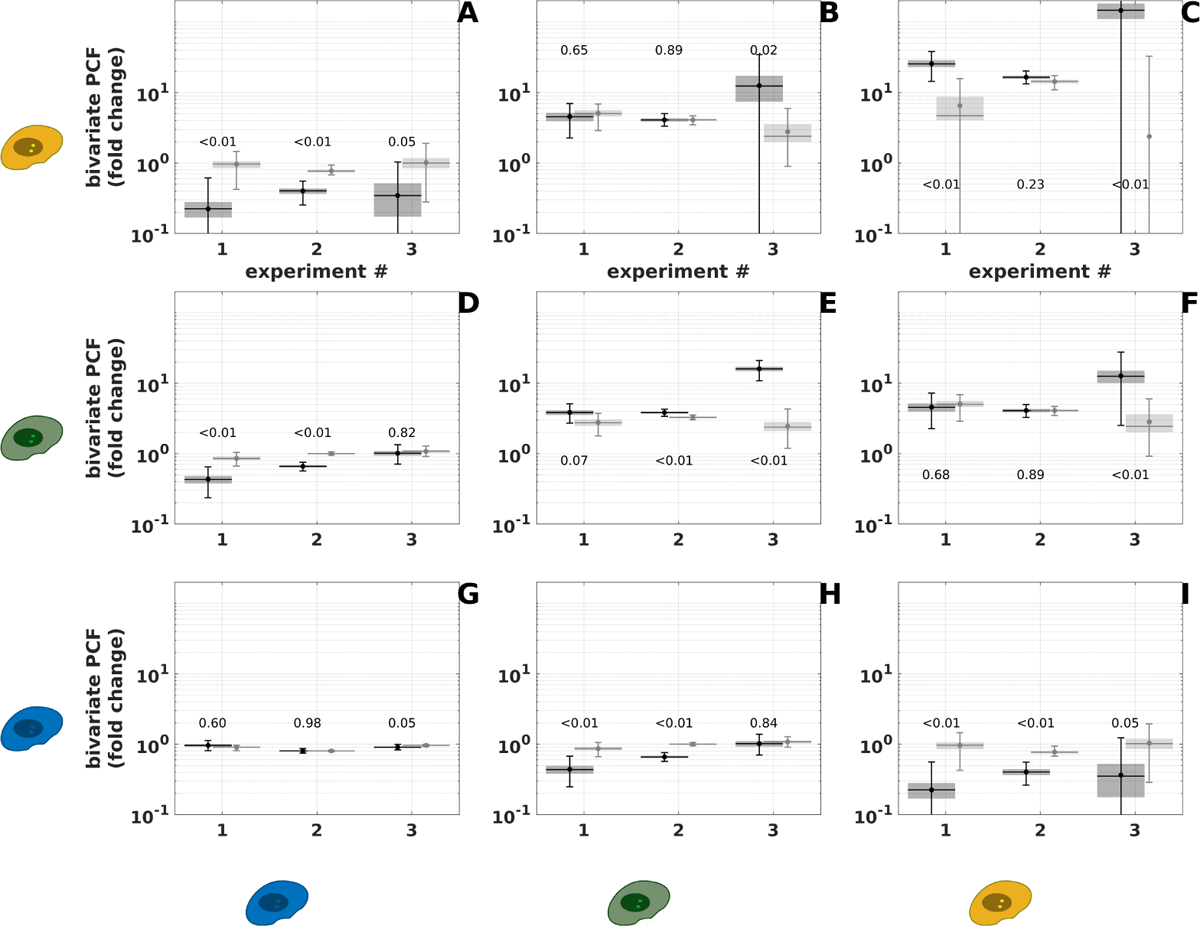
Repeatability of spatio-temporal simulations with MDA-MB-231 cells. We report the ratio between the measured bivariate PCF and the one obtained for phenotype shuffing (control) for data (black) and simulations (gray). This ratio is estimated at *r* = 15μm for which the shuffling PCF is maximum. Dot: actual ratio. Horizontal line: median obtained from bootstrap resampling. Gray shaded box: 50% confidence interval. Error bars: 99% confidence interval. p-values are shown for each data point. p-values are estimated by 500 repeats of simulations to test the null hypothesis (simulations identical to data within the 68% confidence interval obtained from bootstrap resampling). (**A**) Bivariate PCF measuring spatial correlation between CSC and CDC. (**B**) Bivariate PCF measuring spatial correlation between CSC and iCC. (**C**) Bivariate PCF measuring spatial auto-correlation of CSC. (**D**) Bivariate PCF measuring spatial correlation between iCC and CDC. (**E**) Bivariate PCF measuring spatial auto-correlation between iCC. (**F**) Bivariate PCF measuring spatial correlation between iCC and CSC. (**G**) Bivariate PCF measuring spatial auto-correlation of CDC. (**H**) Bivariate PCF measuring spatial correlation between CDC and iCC. (**I**) Bivariate PCF measuring spatial correlation between CDC and CSC.

**Figure 6—figure supplement 1.**
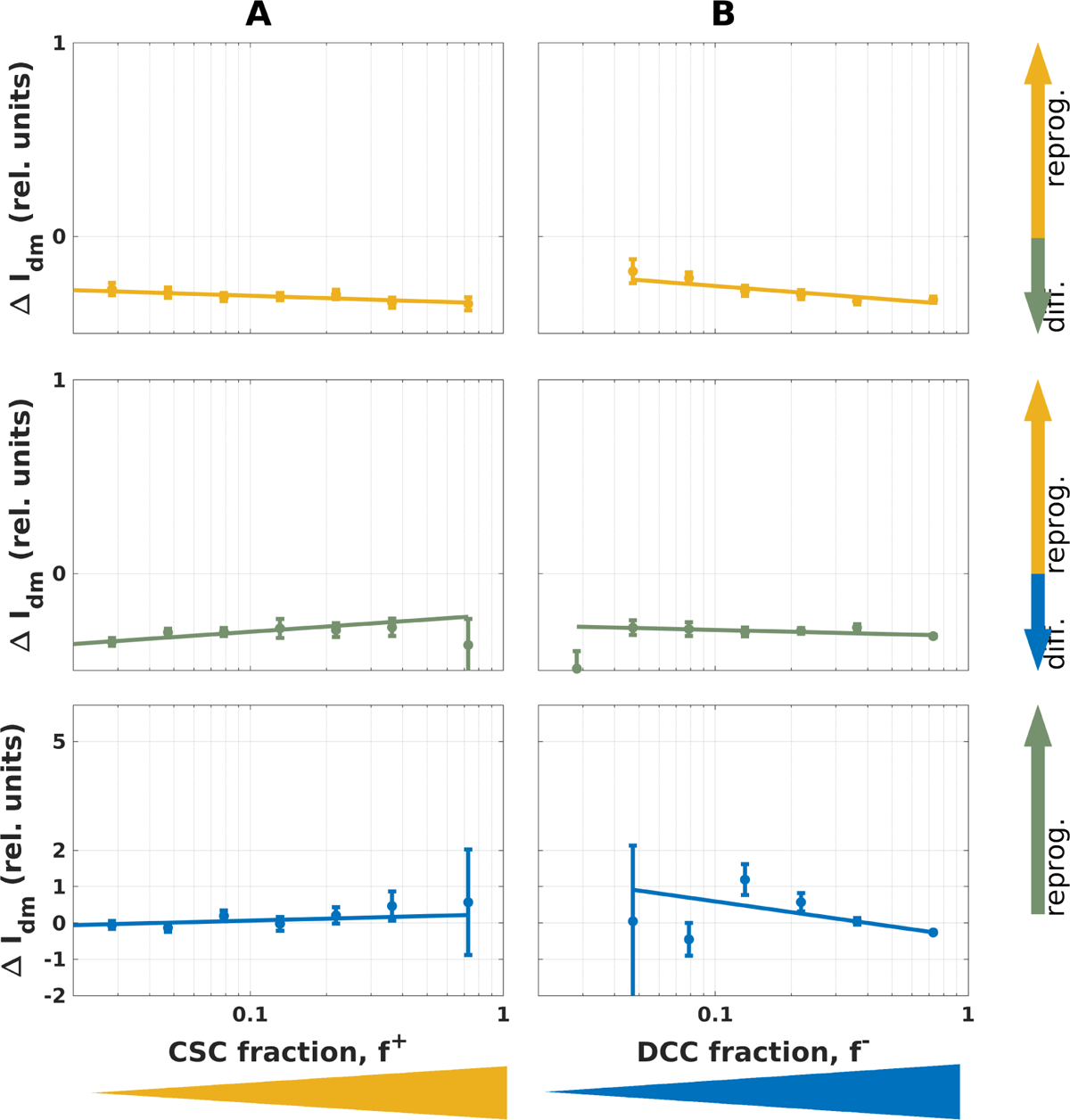
Influence of local environment on fluorescence variation upon mitosis in SUM59PT cells. Average fluorescence variation upon mitosis, ΔI_dm_ conditional to either CSC fraction, f ^+^, (**A**) or CDC fraction, f ^−^, (**B**). From top to bottom data are shown for CSC (yellow), iCC (green) and CDC (blue). ΔI_dm_ is normalized to population average fluorescence intensity of CSC (∼ 150 RFU), iCC (∼ 42 RFU) and CDC (∼ 2.5 RFU). Positive values for ΔI_12_ indicate differentiation and negative values indicate reprogram*min*g. Each point represent conditional mean and the height of error bars two standard deviations.

**Figure 6—figure supplement 2.**
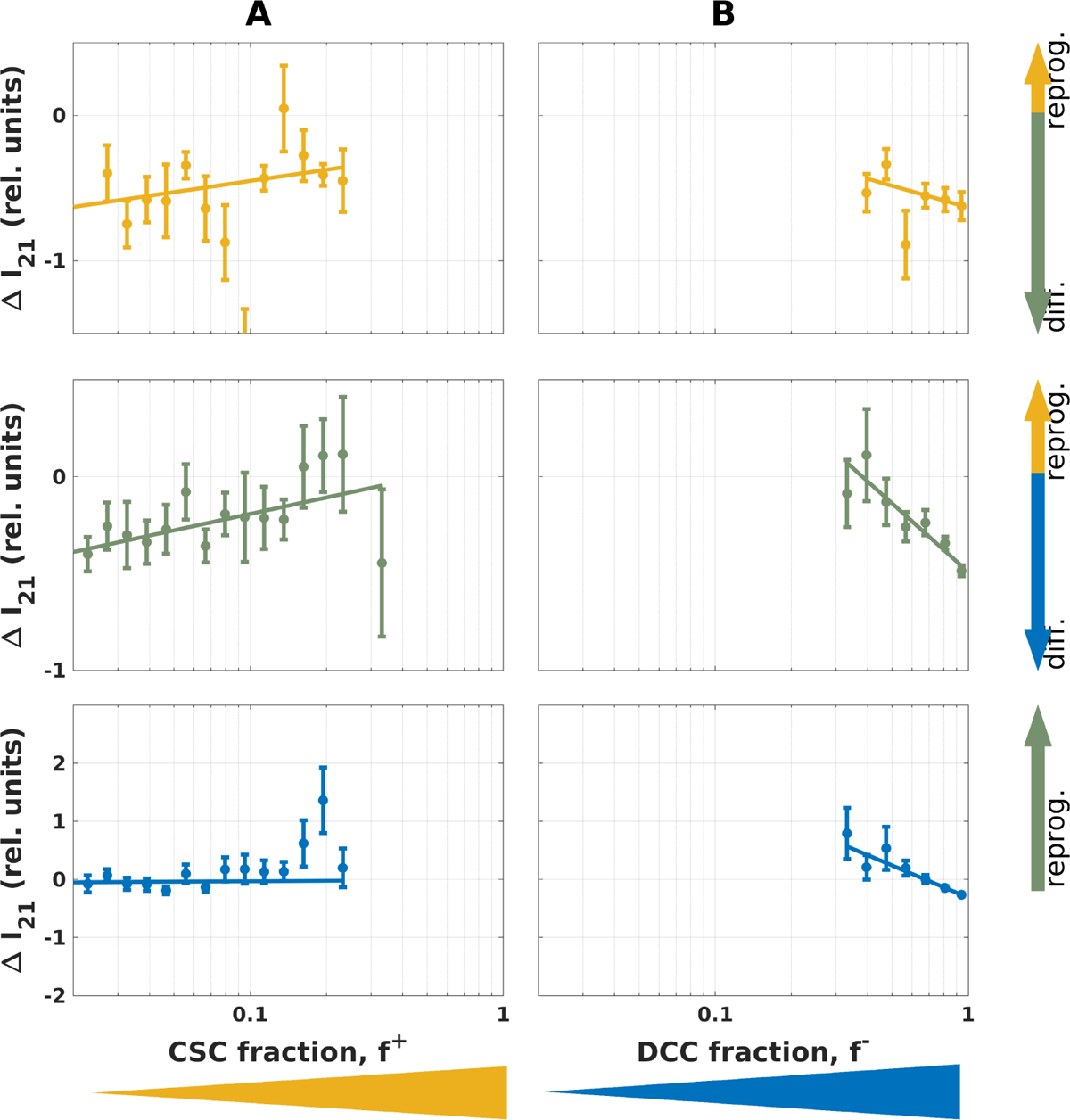
Influence of local environment on fluorescence variation during cell cycle in MDA-MB-231 cells Average fluorescence variation during cell cycle, ΔI_21_ as function of the local CSC fraction, f ^+^, (**A**) or the localCDC fraction, f ^−^, (**B**). From top to bottom data are shown for CSC (yellow), iCC (green) and CDC (blue). ΔI_21_ is normalized to population average fluorescence intensity of CSC (∼ 150 RFU), iCC (∼ 42 RFU) and CDC (∼ 2.5 RFU). Positive values for ΔI_21_ indicate differentiation and negative values indicate reprogram*min*g. Each point represent conditional mean and the height of error bars two standard deviations.

**Figure 6—figure supplement 3.**
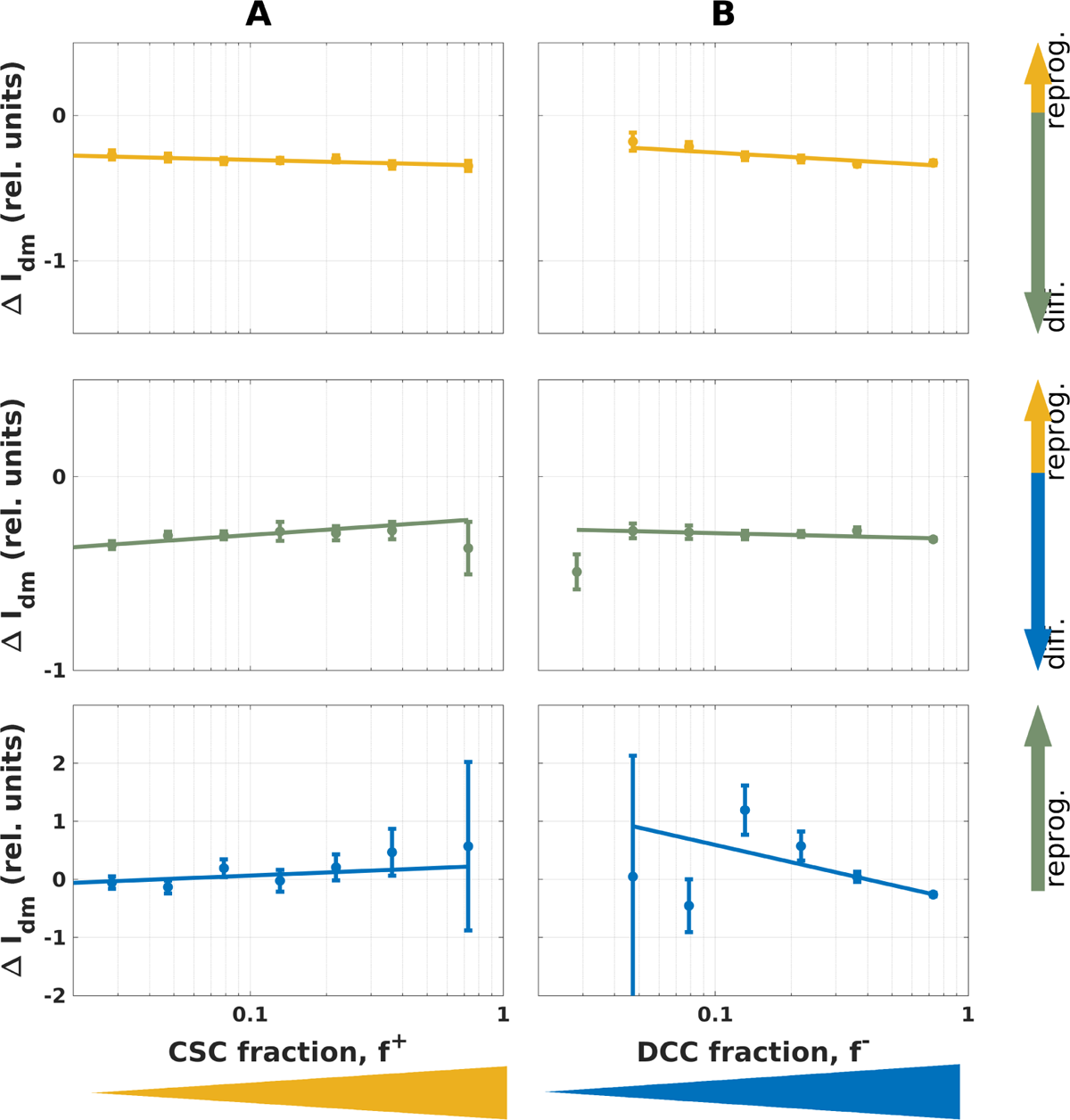
Influence of local environment on fluorescence variation upon mitosis in MDA-MB-231 cells We report average fluorescence variation upon mitosis, ΔI_dm_ conditional to either CSC fraction, f ^+^, (**A**) or CDC fraction, f ^−^, (**B**). From top to bottom data are shown for CSC (yellow), iCC (green) and CDC (blue). ΔI_dm_ is normalized to population average fluorescence intensity of CSC (∼ 150 RFU), iCC (∼ 42 RFU) and CDC (∼ 2.5 RFU). Positive values for ΔI_12_ indicate differentiation and negative values indicate reprogram*min*g. Each point represent conditional mean and the height of error bars two standard deviations.

## Notes

### Competing Interest Statement

The authors have declared no competing interest.

